# Transcription factor clusters enable target search but do not contribute to target gene activation

**DOI:** 10.1101/2022.12.13.520200

**Authors:** Joseph V.W. Meeussen, Wim Pomp, Ineke Brouwer, Wim J. de Jonge, Heta P. Patel, Tineke L. Lenstra

**Affiliations:** Division of Gene Regulation, The Netherlands Cancer Institute, Oncode Institute, 1066CX Amsterdam, the Netherlands

**Keywords:** Gene regulation, transcription factor, clustering, target search, transcription activation, live-cell microscopy

## Abstract

Many transcription factors (TFs) localize in nuclear clusters of locally increased concentrations, but how TF clustering is regulated and how it influences gene expression is not well understood. Here, we use quantitative microscopy in living cells to study the regulation and function of clustering of the budding yeast TF Gal4 in its endogenous context. Our results show that Gal4 cluster formation is facilitated by, but does not completely depend on DNA binding and intrinsically disordered regions. Gal4 cluster properties are regulated by the Gal4-inhibitor Gal80 and Gal4 concentration. Moreover, we discover that clustering acts as a double-edged sword: self-interactions aid TF recruitment to target genes, but recruited Gal4 molecules that are not DNA-bound do not contribute to, and may even inhibit, transcription activation. We propose that cells need to balance the different effects of TF clustering on target search and transcription activation to facilitate proper gene expression.

## INTRODUCTION

Gene-specific transcription factors (TFs) are essential for correct control of gene expression. Eukaryotic TFs contain a DNA-binding domain (DBD) which binds to specific sequences in regulatory promoter and enhancer regions, and a transactivation domain (AD) that interacts with cofactors, chromatin remodelers and other transcriptional regulators to facilitate transcription^1^. Already more than 25 years ago, it was observed that the glucocorticoid receptor TF was not homogeneously distributed through the nucleus, but forms areas of high local concentration, referred to as clusters, hubs, condensates or droplets^2^. These clusters have since been observed for many other TFs, cofactors and RNA polymerase II^3–11^. Despite the widespread observation of clustering, the regulation and function of TF clustering is not well understood.

It has been suggested that cluster formation is driven by multivalent interactions between intrinsically disordered regions (IDRs)^4,12,13^. IDRs are enriched in the transactivation domains of many TFs and enable self-interactions (homotypic interactions) and multivalent interactions with IDRs of other components of the transcriptional machinery (heterotypic interactions)^4,11,12,14–16^. Support for the role of IDRs in TF cluster formation comes from the finding that interactions between the IDRs of TFs and Mediator are important for TF cluster formation *in vitro*^4,17^. However, clustering *in vivo* can also occur independently of IDRs, such as for Sox2 and the glucocorticoid receptor^18,19^. For Sox2, clustering is mostly dependent on DNA binding^18^, suggesting that these clusters reflect binding to adjacent motifs in the genome rather than protein-protein interactions. How endogenous TF clusters are regulated by IDRs, the configuration of binding sites and interactions with DNA or other regulators is only starting to emerge^11,17,20^.

Moreover, an important open question is how clustering influences TFs during the different steps of transcription activation^13,21–23^. Clustering and IDR-mediated interactions have been reported to enhance target search (increasing the DNA binding rate)^24–28^, to increase the local concentration of TFs at the promoter (increasing the DNA binding rate), to stabilize TF binding to DNA (decreasing the rate of TF unbinding)^12^, to enable 3D genomic interactions between target genes^29^ and to boost transcription activation through enhanced recruitment of cofactors and polymerase molecules^30,20,16,31,32^. In contrast, clustering of synthetic TFs and the oncogenic TF EWS::FLI1 instead inhibit gene expression^33,34^. These discrepancies illustrate our lack of understanding of how clustering impacts transcription. Although novel inducible artificial clustering tools provide precise control of clustering, it remains unclear how these results can be extrapolated to endogenous gene regulation.

Here, we used the transcription factor Gal4 from budding yeast to study the regulation and function of TF clustering in an endogenous context. This inducible TF is tightly regulated by the availability of different carbon sources^35^. In cells grown in raffinose, Gal4 is expressed, but is kept inactive by the inhibitor Gal80^36^. In the presence of galactose, Gal80-mediated repression is relieved and Gal4 activates the expression of the *GAL* genes to metabolize galactose. Additionally, expression of *GAL4* as well as its target genes is repressed in the presence of glucose by Mig1 in a concentration dependent manner^37^. The naturally low protein levels of Gal4 and the small number of Gal4 target genes make Gal4 an excellent model to study the effects of TF clustering on transcription at a single locus in an endogenous context.

Using quantitative live-cell imaging of Gal4-EGFP, we find that Gal4 forms clusters *in vivo* that colocalize with target genes. Gal4 cluster abundance, size and density change across different growth conditions, are dependent on the Gal4 expression levels and are limited by interactions with the inhibitor Gal80. Removal of endogenous Gal4 binding sites and analysis of truncation mutants showed that both DNA binding and IDRs contribute to, but are not essential for Gal4 clustering. In addition, regions outside of the Gal4 DNA binding domain are sufficient to recruit additional Gal4 molecules to clusters at target genes, indicating that self-interactions between Gal4 molecules facilitate target search. However, non-DNA-bound Gal4 molecules present in a cluster at a target locus do not necessarily contribute to transcription and might even inhibit transcription. Taken together, we propose that clustering positively affects target search and negatively affects transcription activation, and these aspects therefore need to be properly balanced to facilitate gene expression.

## RESULTS

### Gal4 forms clusters in living yeast cells

To visualize endogenous Gal4 in living yeast cells, we fused EGFP with a flexible linker to the C-terminus of Gal4. Addition of the EGFP tag did not affect cell growth on galactose-containing plates, indicating full functionality in inducing the *GAL* genes (Figure S2A). *GAL4-EGFP* cells were grown in media with different carbon sources, where the *GAL* genes are either repressed (glucose), uninduced (raffinose) or induced (galactose)^38^, and imaged in 3D with high signal-to-noise ratio using highly inclined and laminated optical sheet (HILO) microscopy^39^. In all three growth conditions, we observed bright Gal4-EGFP foci of high local concentration (Figure 1B), hereafter referred to as Gal4 clusters. These clusters were not observed in the WT strain (without *EGFP*) or a strain expressing nuclear *EGFP* (Figure S2B), indicating that the observed clustering is specific for Gal4. To quantify the size and intensity of the observed clusters, they were fit with a 3D gaussian model. This allowed extraction of the peak width (the standard deviation *σ*, a measure for cluster size) and peak height (cluster density, a measure for Gal4 concentration within the cluster), which together determine the total intensity (the integrated peak intensity, a measure for the total number of molecules in the cluster) (Figure S1). Fitting of beads confirmed the independence of the size and density in the fitting algorithm and allowed us to use *σ* to estimate the cluster diameter (Figure S1). Quantification and localization of the clusters with this algorithm revealed that both the number of Gal4 clusters and the total cluster intensity were lowest in repressed cells (glucose), intermediate in uninduced cells (raffinose) and highest in induced cells (galactose) (Figure 1C, F). Repressed cells showed less dense clusters than uninduced and induced cells, while induced cells showed larger clusters than repressed or uninduced cells (Figure 1D-E). In induced conditions, the cluster diameter ranged approximately between 100 and 750 nm (Figure S1F). We conclude that the TF Gal4 forms clusters and that the degree of clustering positively correlates with conditions of active *GAL* gene transcription.

**Figure 1:**
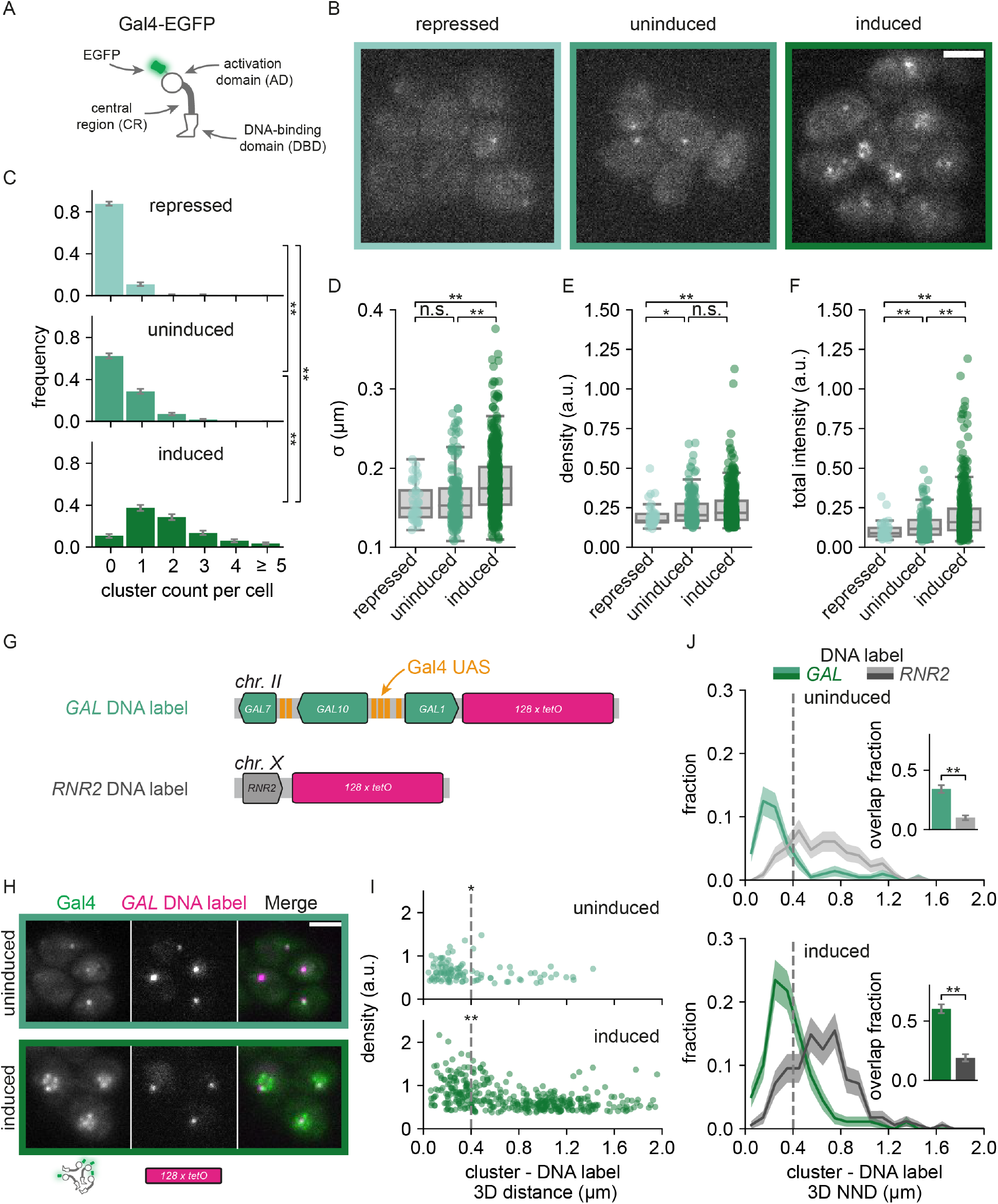
Gal4 forms clusters that colocalize with the *GAL* genes. **A.** Schematic representation of Gal4. The activation domain (AD), central region (CR) and DNA-binding domain (DBD) are indicated. For visualization, the C-terminus of the endogenous Gal4 is tagged with EGFP (green). Note that Gal4 binds DNA as a dimer. **B.** Representative images of Gal4-EGFP clusters in yeast cells grown in repressed (glucose, very light green), uninduced (raffinose, light green) and induced (galactose, green) conditions. Images are a single *z*-slice of a representative group of cells. Scalebar: 3 μm. **C-F.** Quantification of Gal4-EGFP clusters in repressed (very light green, 285 cells), uninduced (light green, 381 cells) and induced (dark green, 280 cells) conditions. **C.** Distribution of number of clusters observed per cell. Error bars indicate standard error of the mean (SEM) based on 1000 bootstrap repeats. **D-F.** Distribution of **D.** cluster *σ*, **E.** density and **F.** total intensity, represented by the sigma, peak height and integrated intensity of the 3D gaussian fit, respectively (see methods for details). Circles show data for individual clusters and box plots show the distribution of the data, with box edges indicating first and third quartiles, center line indicating the median and whiskers indicating the 1.5× interquartile range. Significance determined by Mann-Whitney *U* test; n.s.: not significant; *: p < 0.05; **: p < 0.01. **G.** Schematic representation of genomic integration of the *128xtetO* DNA label (magenta) at the *GAL* locus (light green), with Gal4 bindings sites (UAS, orange), or at the *RNR2* locus (grey), without Gal4 binding sites. **H.** Representative images of dual-color fluorescence imaging to determine the colocalization between Gal4-EGFP clusters (green) and the *GAL* locus (magenta) in uninduced (raffinose) and induced (galactose and raffinose) conditions. Images are a single *z*-slice of a representative group of cells. Scalebar: 3 μm. **I.** Scatterplot of Gal4-EGFP cluster density versus 3D distance between the cluster and the *GAL* DNA label for uninduced (top, dark green) and induced (bottom, light green) conditions (306 and 206 cells, respectively). Vertical dashed line indicates 400 nm threshold used to discriminate between overlapping and non-overlapping clusters. Significance between clusters closer and further than 400 nm from the DNA label was determined by Mann-Whitney *U* test; *: p < 0.05; **: p < 0.01. **J.** Distribution of 3D nearest neighbor distances (NND) between the *GAL* (green) and *RNR2* (grey) DNA label and the closest Gal4-EGFP cluster in uninduced (top, light green and light grey, 306 and 276 cells, respectively) and induced conditions (bottom, dark grey and dark green, 200 and 198 cells, respectively). Shaded regions represent SEM based on 1000 bootstrap repeats. Vertical dashed line indicates 400 nm threshold used to discriminate between overlapping and non-overlapping clusters. Inset shows fraction of *GAL* or *RNR2* loci with an overlapping cluster. Error bars represent SEM based on 1000 bootstrap repeats. Significance determined by Fisher’s exact test; n.s.: not significant; **: p < 0.01.

### Gal4 clusters are enriched at their endogenous target genes

To test whether these Gal4 clusters localize at endogenous target sites, we integrated 128 repeats of the *tetO* sequence downstream of the *GAL1* gene and constitutively expressed tetR-tdTomato to visualize the location of the *GAL1-GAL10-GAL7* locus inside living cells (Figure 1G)^40^. The *GAL* locus contains six Gal4 binding sites: four in the *GAL1-10* promoter and two in the *GAL7* promoter. Dual-color imaging of Gal4-EGFP clusters and the *GAL* DNA label in uninduced and induced conditions revealed frequent colocalization of Gal4 clusters at the *GAL* genes (Figure 1H). In both conditions the densest Gal4 clusters were found in proximity (3D distance < 400 nm) of the *GAL* locus (Figure 1I). These close-proximity clusters were, however, slightly smaller than clusters further away from the locus (Figure S2E), such that the total intensity was increased but not as much as the density (Figure S2F). Binding to the multiple UASs in the *GAL* locus may thus result in more concentrated, smaller Gal4 clusters.

To quantify the colocalization of clusters with the *GAL* locus, we determined the 3D nearest neighbor distance (NND) from each DNA label to a Gal4 cluster, which showed a clear peak at proximal distances (Figure 1I). To discriminate clusters that overlap with the *GAL* locus, the percent overlap was calculated using different distance thresholds (Figure S2D). We chose a threshold of 400 nm to define overlapping clusters, since this threshold included the majority of the proximal high-intense clusters in uninduced conditions, while simultaneously limiting random overlap within the small yeast nucleus (< 2 μm diameter). We note that the NND distribution is wider in induced than in uninduced condition (Figure 1I), perhaps because the *GAL* locus expands upon transcription activation, and that the 400 nm threshold likely results in an underestimate of the true overlap. Throughout the manuscript, we ensured that the results are independent of the chosen threshold. Using a threshold of 400 nm, 34 ± 3% and 60 ± 4% of the *GAL* loci overlap with a Gal4 cluster in uninduced and induced cells, respectively (Figure 1J, inset).

To check whether this colocalization was specific for the *GAL* genes, the DNA label was also placed in a different yeast strain downstream (3’) of *RNR2*, a gene located on a chromosome without any *GAL* genes and that transcribes independently of Gal4 binding^41^ (Figure 1G, J). In contrast to the *GAL* DNA label, the brightest Gal4 clusters were not enriched at the *RNR2* 3’ DNA label (Figure 1J). In addition, compared to the *GAL* genes, the distribution of NNDs at *RNR2* was much broader (Figure 1J) and only showed 10 ± 2% and 19 ± 3% overlap in uninduced and induced conditions, respectively. A second version of the *RNR2* DNA label, in which the label was placed upstream (5’) of *RNR2*, showed similar results (Figure S2G-H). To confirm these results and to eliminate a possible bias from the lower *z*-resolution compared to the *x-y* resolution, we repeated this analysis in 2D (*xy*), which also showed many more proximal Gal4 clusters at the *GAL* genes than at the *RNR2* gene (Fig S2I). These results indicate that Gal4 clusters are enriched at the *GAL* locus in both uninduced and induced conditions.

### The repressor Gal80 limits Gal4 cluster formation

As Gal4 clustering differs between growth conditions, we investigated how Gal4 clustering is regulated. We focused on regulatory features that differ between different sugar conditions, including Gal80-mediated repression of Gal4 activity, Gal4 interactions with the transcriptional machinery and Gal4 expression levels. In repressed and uninduced conditions (glucose and raffinose), Gal80 represses Gal4 activity. To test whether Gal80 repression limits Gal4 clustering in uninduced conditions, we analyzed Gal4 clustering in a *gal80*Δ strain in raffinose (Figure 2A). Western blot analysis of Gal4-EGFP-V5 verified that the Gal4 expression levels in *gal80*Δ cells were comparable to WT *GAL80* cells (Figure S3B). Deletion of *GAL80* increased Gal4 cluster abundance, cluster size, cluster density and total cluster intensity (Figure 2B-E), indicating that Gal80 suppresses the clustering capability of Gal4 in uninduced conditions.

**Figure 2:**
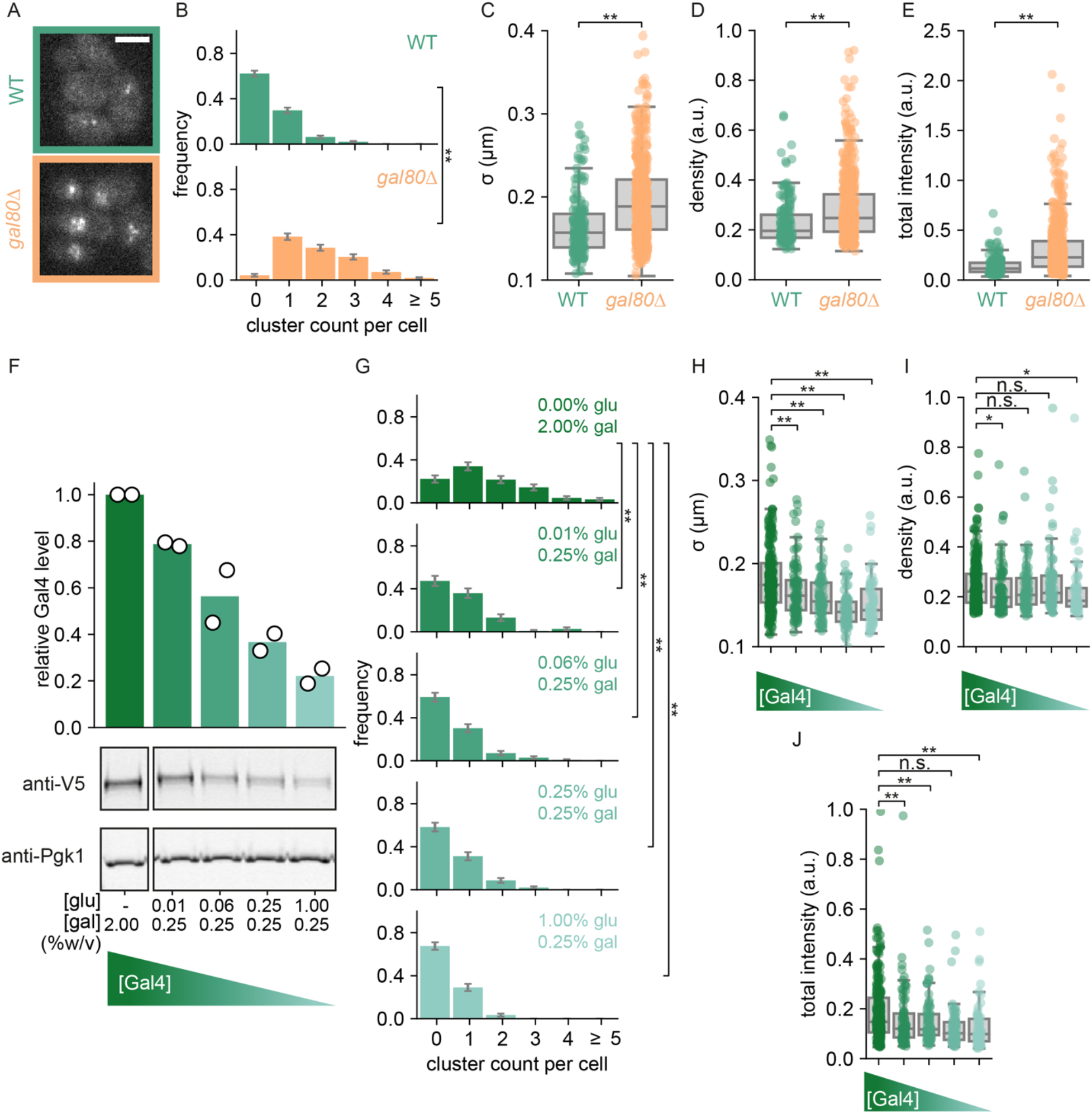
Gal4 cluster formation is inhibited by Gal80 and dependent on Gal4 concentration. **A.** Representative images of Gal4-EGFP clusters in WT (green) and *gal80*Δ (orange) yeast cells in uninduced (raffinose) conditions. Images are a single *z*-slice of a representative group of cells. Scalebar: 3 μm. **B-E.** Quantification of Gal4-EGFP clusters in WT (green, 357 cells) and *gal80*Δ (orange, 285 cells) in uninduced (raffinose) conditions. **B.** Distribution of number of clusters observed per cell. Error bars indicate SEM based on 1000 bootstrap repeats. **C-E.** Distribution of **C.** cluster *σ*, **D.** density and **E.** total intensity. Circles show data for individual clusters and box plots show the distribution of the data, with box edges indicating first and third quartiles, center line indicating the median and whiskers indicating the 1.5× interquartile range. Significance determined by Mann-Whitney *U* test; **: p < 0.01. **F.** Western blot quantification of Gal4-EGFP-V5 protein levels using an anti-V5 antibody measured in the *gal80*Δ background across a range of glucose and galactose concentrations (%w/v) as indicated. Expression levels are normalized to Pgk1 and to the expression level in 2% galactose in the absence of glucose. Open circles represent the results of individual replicate experiments, bars indicate their mean. Western blot images are a representative example of 2 independent experiments. **G-J.** Quantification of Gal4-EGFP clusters in *gal80*Δ background in the same glucose and galactose conditions as in F., with in total 153, 114, 142, 151 and 179 cells included in the respective conditions. G. Distribution of number of clusters observed per cell. Error bars indicate SEM based on 1000 bootstrap repeats. H-J. Distribution of H. cluster s, I. density and J. total intensity. Circles show data for individual clusters and box plots show the distribution of the data, with box edges indicating first and third quartiles, center line indicating the median and whiskers indicating the 1.5× interquartile range. Significance determined by Mann-Whitney *U* test; n.s.: not significant; *: p < 0.05; **: p < 0.01.

### Gal4 interactions with the Mediator tail are not required for Gal4 clustering

When Gal80 inhibition is relieved, Gal4 interacts with Mediator via dynamic “fuzzy” interactions with the Med15 subunit^15^. Med15 forms phase-separated condensates *in vivo* in mammalian cells^42^, and Mediator condensates are able to induce phase-separation of ADs of various TFs *in vitro*^4^. Because Gal80 shields the Gal4 AD and limits Gal4 clustering, we tested whether interactions of Gal4 with Med15 are important for Gal4 clustering. Comparison of Gal4 clusters between WT and *med15*Δ cells revealed that clusters were slightly more abundant, more intense and larger upon Med15 deletion (Figure S3A, C-E). These results indicate that interactions with Med15 are not required for Gal4 clustering and may even limit them. Inhibition of Gal4 cluster formation by Gal80 thus occurs independent of suppression of Gal4-Mediator interactions. Gal80 may instead suppress Gal4-Gal4 self-interactions by physically shielding the AD or by enabling a more structured conformation of the AD^15^.

### Gal4 clustering is concentration-dependent

In addition to Gal80-mediated regulation of Gal4 activity, *GAL4* expression is repressed by Mig1 in the presence of glucose^37^. In line with previous studies, cells grown in repressed conditions showed reduced Gal4-EGFP-V5 levels by western blot compared to uninduced and induced conditions (Figure S2C). Surprisingly, however, Gal4-EGFP-V5 levels were higher in cells grown in galactose than in raffinose, even though galactose was previously shown not to increase Gal4 mRNA levels, nor increase expression of a reporter gene driven by the *GAL4* promoter^43,44^. We speculate that the Gal4 protein level increase in galactose may be caused by changes in its protein degradation rate. Regardless, these differences in Gal4 protein levels across these sugar conditions raised the question whether Gal4 clustering is concentration dependent, as has been found for other TFs^4,30,34^.

To test the relationship between Gal4 concentration and Gal4 clustering, we varied the Gal4 concentration by modulating the glucose concentration (Figure 2F)^37^ and analyzed Gal4 clustering at these varying Gal4 protein levels (Figure 2G-J). These experiments were performed in *gal80*Δ cells to prevent confounding effects of Gal80-regulation on Gal4 clusters. At higher Gal4 protein levels, more and larger clusters were observed, with minor differences in cluster density (Figure 2G-J). Overall, we conclude that Gal4 clusters are regulated by Gal80 and are concentration dependent.

### DNA binding facilitates, but is not essential for Gal4 clustering

Next, we examined how Gal4 clustering is affected by DNA binding. Gal4 binds to a 17-bp 5′-CGG-N11-CCG-3′ consensus sequence (upstream activating sequence, UAS) as a dimer^45,46^. Since the promoters of several Gal4 target genes contain multiple Gal4 UASs: 4 UASs in *pGAL1-10*, 2 UASs in *pGAL7* and 2 UASs in *pGAL2* (Figure 3A), clusters could possibly arise from binding of several Gal4 molecules to adjacent motifs in these promoters. We tested this by scrambling all-but-one UAS for each of these three target promoters, such that the consensus sequence is lost (scrUAS, Figure 3A) and no adjacent Gal4 bindings sites are left in the genome^47^. In this scrUAS strain, Gal4 clusters do not disappear and were only mildly affected (Figure 3B-F), indicating that Gal4 clusters do not simply reflect multiple Gal4 molecules that are bound to adjacent binding sites. When adjacent UASs are scrambled, cluster abundance is only slightly reduced compared to a WT UAS stain, but interestingly, the clusters become larger and less dense (Figure 3E-F). Binding to adjacent sites, in particular at the *GAL1-GAL10-GAL7* locus, may thus enhance cluster concentration and reduce their size (Figure 3D-E).

**Figure 3:**
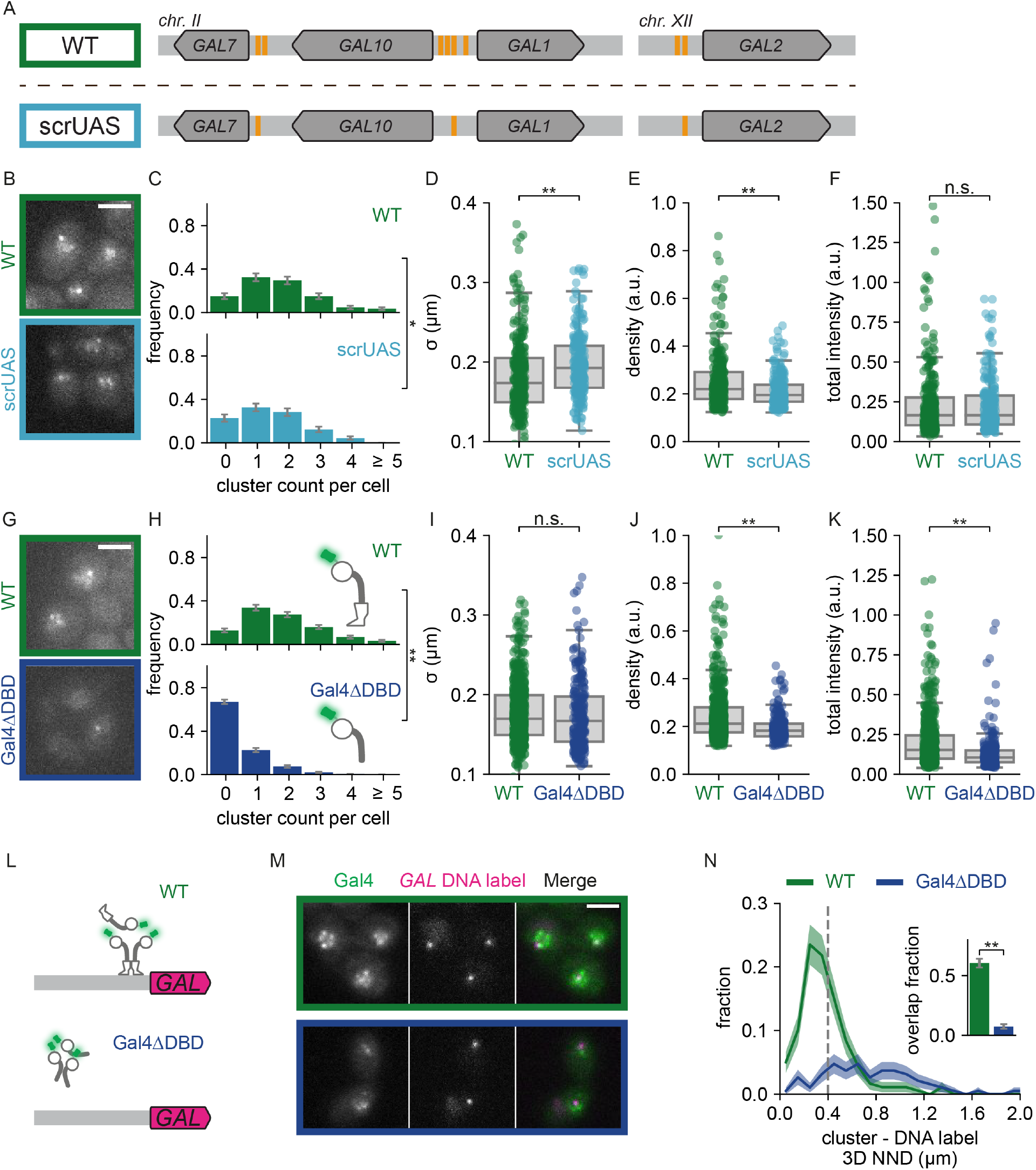
DNA binding is not essential for Gal4 clustering but is essential for cluster localization. **A.** Schematic representation of the *GAL* genes with multiple UAS sites (orange) in WT (top, green box) and in a mutant with scrambled UAS sites (scrUAS; bottom, cyan box) such that only a single UAS site remains present in the promoter of each *GAL* gene (*scrUAS 5’GAL2* + *scrUAS 5’GAL7* + 3× *scrUAS 5’GAL10*, bottom, cyan box). **B.** Representative images of Gal4-EGFP clusters in WT (green) and scrUAS (cyan) yeast cells in induced (galactose and raffinose) conditions. Images are a single *z*-slice of a representative group of cells. Scalebar: 3 μm. **C-F.** Quantification of Gal4-EGFP clusters in WT (green, 173 cells) and scrUAS (orange, 163 cells) in induced (galactose and raffinose) conditions. **C.** Distribution of number of clusters observed per cell. Error bars indicate SEM based on 1000 bootstrap repeats. **D-F.** Distribution of **D.** cluster *s*, **E.** density and **F.** total intensity. Circles show data for individual clusters and box plots show the distribution of the data, with box edges indicating first and third quartiles, center line indicating the median and whiskers indicating the 1.5× interquartile range. Significance determined by Mann-Whitney *U* test; n.s.: not significant; *: p < 0.05; **: p < 0.01. **G.** Representative images of WT (Gal4-EGFP; green) and Gal4ΔDBD (BPSV40-Gal4(Δ2-94)-EGFP; blue) clusters in yeast cells in induced (galactose and raffinose) conditions. Images are a single *z*-slice of a representative group of cells. Scalebar: 3 μm. **H-K.** Same as **C-F.** for WT Gal4 (Gal4-EGFP; green, 313 cells) and Gal4ΔDBD (BPSV40-Gal4(Δ2-94)-EGFP; blue, 517 cells) clusters in induced conditions. Significance determined by Mann-Whitney *U* test; n.s.: not significant; **: p < 0.01. **L.** Schematic representation of the *GAL* locus with DNA label (magenta) and clusters of either WT (Gal4-EGFP) or Gal4ΔDBD (Gal4ΔDBD-EGFP). **M.** Representative images of dual-color fluorescence imaging to determine the colocalization between clusters (green) of either WT (green outline) or Gal4ΔDBD (blue outline) and the *GAL* locus (magenta) in induced conditions. Images are a single *z*-slice of a representative group of cells. Scalebar: 3 μm. **N.** Distribution of 3D nearest neighbor distances (NND) between the *GAL* DNA label and the closest cluster of WT Gal4 (green, 200 cells) and Gal4*Δ*DBD (blue, 211 cells). Shaded regions represent SEM based on 1000 bootstrap repeats. Vertical dashed line indicates 400 nm threshold used to discriminate between overlapping and non-overlapping clusters. Inset shows fraction of *GAL* loci with an overlapping cluster. Error bars represent SEM based on 1000 bootstrap repeats. Significance determined by Fisher’s exact test; **: p < 0.01.

To investigate further how DNA binding affects Gal4 clustering, we deleted the N-terminal region containing both the DNA binding domain (DBD) and the dimerization domain^46,48,49^. As this region also contains the nuclear localization signal (NLS)^50,51^, nuclear localization was ensured by addition of a strong BPSV40 NLS^52^. As expected, the Gal4ΔDBD mutant strain is unable to grow on galactose-containing plates (Figure S4A). We found that deletion of the DBD did not abolish clustering but decreased both cluster abundance and intensity (Figure 3E-I). This reduction in clustering was not caused by lower Gal4 levels, as V5-tagged Gal4ΔDBD levels by western blot were modestly increased compared to Gal4 WT (Figure S4B). In addition, to distinguish between the effect of DNA binding and dimerization, we mutated a single amino acid, S41D, which abolishes Gal4 binding to the endogenous *GAL1/10* promoter *in vivo* and to DNA *in vitro* but still contains intact dimerization^53^. Similar to Gal4ΔDBD, Gal4(S41D) clusters were less abundant and less dense than WT (Figure S4D-G). DNA binding thus contributes to cluster formation and allows for more concentrated clusters, but it is not essential for clustering.

Analysis of Gal4ΔDBD cluster localization using the *GAL* DNA label showed that deletion of the DNA binding domain resulted in loss of cluster enrichment (7 ± 2% overlap) at the *GAL* locus, in line with the essential function of DNA binding domains in determining sequence-specificity (Figure 3G-N, Figure S4H). Together, these results indicate that in the absence of DNA binding clusters can no longer localize at target genes, but can still form, albeit at a much-reduced rate.

### IDRs are not essential for Gal4 clustering, but contribute to target search

Our results show that the DBD of Gal4 is essential for cluster localization to the *GAL* genes, which is consistent with the sequence-specificity of DBDs. In addition, several reports have indicated that TF target gene selection is facilitated by regions outside DBDs^24,54,55^. In some cases, the DBD is even dispensable for target search^24,56^. For Gal4, the central region (CR) and C-terminal AD enhance localization of the DBD to an *in vivo* reporter array^57^. Both these regions have been predicted and shown to contain disordered regions^15,58,59^ (Figure S5A). Since IDRs have previously been linked to cluster formation and target search^13,24,25^, we wondered how the CR and AD of Gal4 contribute to cluster formation and localization.

To address this, we constructed three truncation mutants: (1) Gal4ΔminiAD, lacking the last 40 amino acids of the AD; (2) Gal4ΔAD, lacking the entire disordered AD; and (3) Gal4-DBD-only, lacking the CR and the AD, and thus consisting only of the DBD and dimerization domain. All Gal4 truncation mutants showed higher Gal4 protein levels compared to WT Gal4 (Figure S4C), and were unable to grow on galactose (Figure S4A). The Gal4ΔminiAD and Gal4ΔAD showed similar or slightly fewer clusters of similar size and density compared to WT (Figure 4A-J), indicating that interactions between the AD and the transcriptional machinery do not have a major effect on clustering. Unexpectedly, the Gal4-DBD-only still formed clusters, despite lacking most, if not all, of the IDRs (Figure 4K). However, clusters were severely reduced in number, size, density and total intensity compared to WT Gal4 (Figure 4L-O). These remaining clusters cannot be explained as binding of multiple Gal4-DBD-only dimers to adjacent UASs, since these Gal4-DBD-only clusters still persisted when all adjacent UASs were scrambled (Figure S5B-E). Together, these truncation experiments demonstrate that Gal4 clustering is mediated by multiple protein domains and that IDRs contribute to, but are not essential for Gal4 clustering.

**Figure 4:**
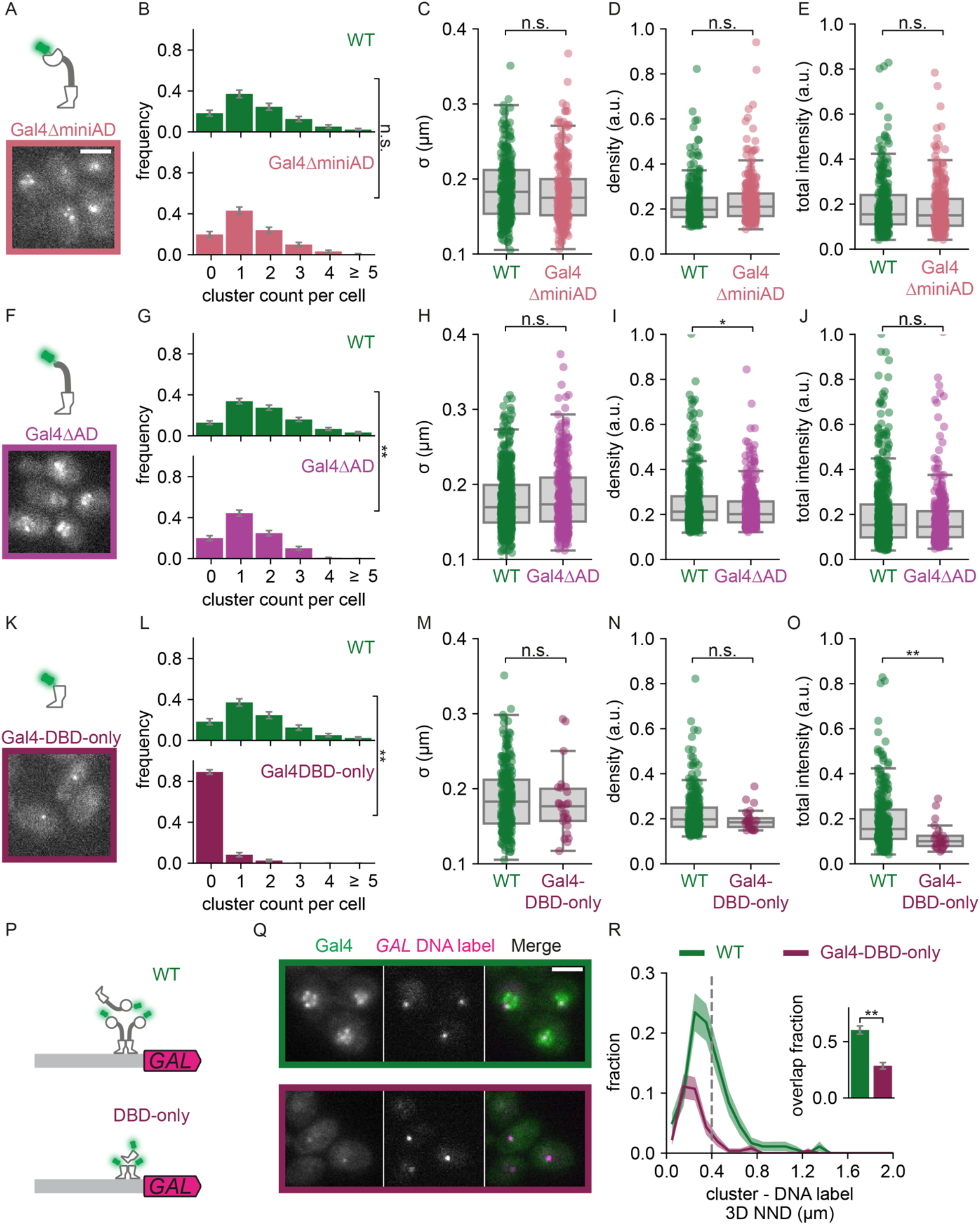
IDRs are not essential for Gal4 clustering but contribute to target search. **A.** Top: Schematic representation of Gal4*Δ*miniAD (Gal4(Δ840-881)-EGFP). Bottom: Representative image of Gal4*Δ*miniAD clusters in yeast cells in induced (galactose and raffinose) conditions. Image is a single *z*-slice of a representative group of cells. **B-E.** Quantification of WT Gal4 (Gal4-EGFP; green, 175 cells) and Gal4ΔminiAD (pink, 175 cells) clusters in induced conditions. **B.** Distribution of number of clusters observed per cell. Error bars indicate SEM based on 1000 bootstrap repeats. **C-E.** Distribution of **C.** cluster *σ*, **D.** density and **E.** total intensity. Circles show data for individual clusters and box plots show the distribution of the data, with box edges indicating first and third quartiles, center line indicating the median and whiskers indicating the 1.5× interquartile range. Significance determined by Mann-Whitney *U* test; n.s.: not significant. **F.** Top: Schematic representation of Gal4*Δ*AD (Gal4(Δ768-881)-EGFP). Bottom: Representative image of Gal4ΔAD clusters in yeast cells in induced conditions. Image is a single *z*-slice of a representative group of cells. **G-J.** Same as **B-E.** for WT Gal4 (green, 313 cells) and Gal4ΔAD (magenta, 270 cells) clusters in induced conditions. Significance determined by Mann-Whitney *U* test; n.s.: not significant; *: p < 0.05; **: p < 0.01. **K.** Top: Schematic representation of Gal4-DBD-only (Gal4(Δ95-881)-EGFP). Bottom: Representative image of Gal4-DBD-only clusters in yeast cells in induced conditions. Image is a single *z*-slice of a representative group of cells. **L-O.** Same as **B-E.** WT Gal4 (green, 175 cells) and Gal4-DBD-only (purple, 193 cells) clusters in induced conditions. Significance determined by Mann-Whitney *U* test; n.s.: not significant; **: p < 0.01. **P.** Schematic representation of the *GAL* locus with DNA label (magenta) and clusters of either WT Gal4 (green) or Gal4-DBD-only (purple). **Q.** Representative images of dual-color fluorescence imaging to determine the colocalization between clusters (green) of either WT Gal4 (green outline) or Gal4-DBD-only (purple outline) and the *GAL* DNA label (magenta) in induced conditions. Images are a single *z*-slice of a representative group of cells. Scalebar: 3 μm. **R.** Distribution of 3D nearest neighbor distances (NND) between the *GAL* DNA label and the closest cluster of WT Gal4 (green, 200 cells) and Gal4-DBD-only (purple, 276 cells). Shaded regions represent SEM based on 1000 bootstrap repeats. Vertical dashed line indicates 400 nm threshold used to discriminate between overlapping and non-overlapping clusters. Inset shows fraction of DNA label-containing cells with an overlapping cluster. Error bars represent SEM based on 1000 bootstrap repeats. Significance determined by Fisher’s exact test; **: p < 0.01.

The large reduction of clustering in the Gal4-DBD-only mutant suggests that the CR and AD contribute to self-interactions that enable cluster formation. We next asked whether these CR and AD-mediated self-interactions contribute to localization of clusters to target genes. To test this, we measured the overlap of the Gal4-DBD-only clusters with the endogenous *GAL* gene locus and found 29 ± 3% overlap (Figure 4P-R). Importantly, in the Gal4-DBD-only mutant many cells did not contain any clusters (Figure 4L, S5F), reducing the number of cells with a Gal4-DBD-only cluster at the *GAL* locus compared to WT (Figure 4R). The IDRs in the CR and AD thus contribute to the Gal4 cluster formation and localization at a target locus, in agreement with previous observations at artificial arrays^57^.

### Gal4 self-interactions are sufficient to recruit Gal4 to target genes

Our mutant analysis indicated that although the CR and ADs of Gal4 facilitate Gal4 clustering (Figure 4), proper localization to the *GAL* locus requires a functional DBD (Figure 3). These findings suggested that the DBD may anchor Gal4 at the correct sites, and that CR- and AD-mediated self-interactions facilitate recruitment of additional Gal4 molecules and cluster formation. To test this model, we assessed whether the GalΔDBD mutant could be recruited to the *GAL* locus through clustering with Gal4 molecules that do contain a DBD. This experiment was performed in a diploid yeast strain where one allele expressed Gal4ΔDBD-EGFP and the other allele contained either a *GAL4*-deletion (*gal4*Δ), full length WT Gal4 or Gal4ΔAD (containing the DBD and CR) (Figure 5A). As expected, in absence of an additional Gal4 copy, the Gal4ΔDBD clusters did not localize at the *GAL* locus (Figure 5A-B, top panel), and only 7 ± 2% of *GAL* DNA labels overlapped with a Gal4ΔDBD cluster (Figure 5B), which is the same as the overlap in haploid cells (7 ± 2%, Figure 3K). However, upon additional expression of WT Gal4, the Gal4ΔDBD-EGFP clusters showed a clear peak of enrichment at the *GAL* locus, and a significantly increased overlap with the *GAL* locus from 7 ± 2% to 13 ± 2% (Figure 5B-C, top panel). Expression of the Gal4ΔAD mutant also showed the same trend of increasing the localization of the Gal4ΔDBD-EGFP clusters to the *GAL* locus (10 ± 2%) (Figure 5B-C, bottom panel). This increase in overlap was independent of the chosen threshold to determine the overlap (Figure S6A). In addition, analysis of the overlap in 2D instead of 3D revealed even clearer peaks of enrichment of Gal4 clusters close to the *GAL* genes in the presence of the Gal4ΔAD or Gal4 WT, but not when the second allele was deleted (Figure S6A-B). These results indicate that the Gal4ΔDBD mutant can be recruited to its target genes via protein-protein interactions with DBD-containing Gal4.

**Figure 5:**
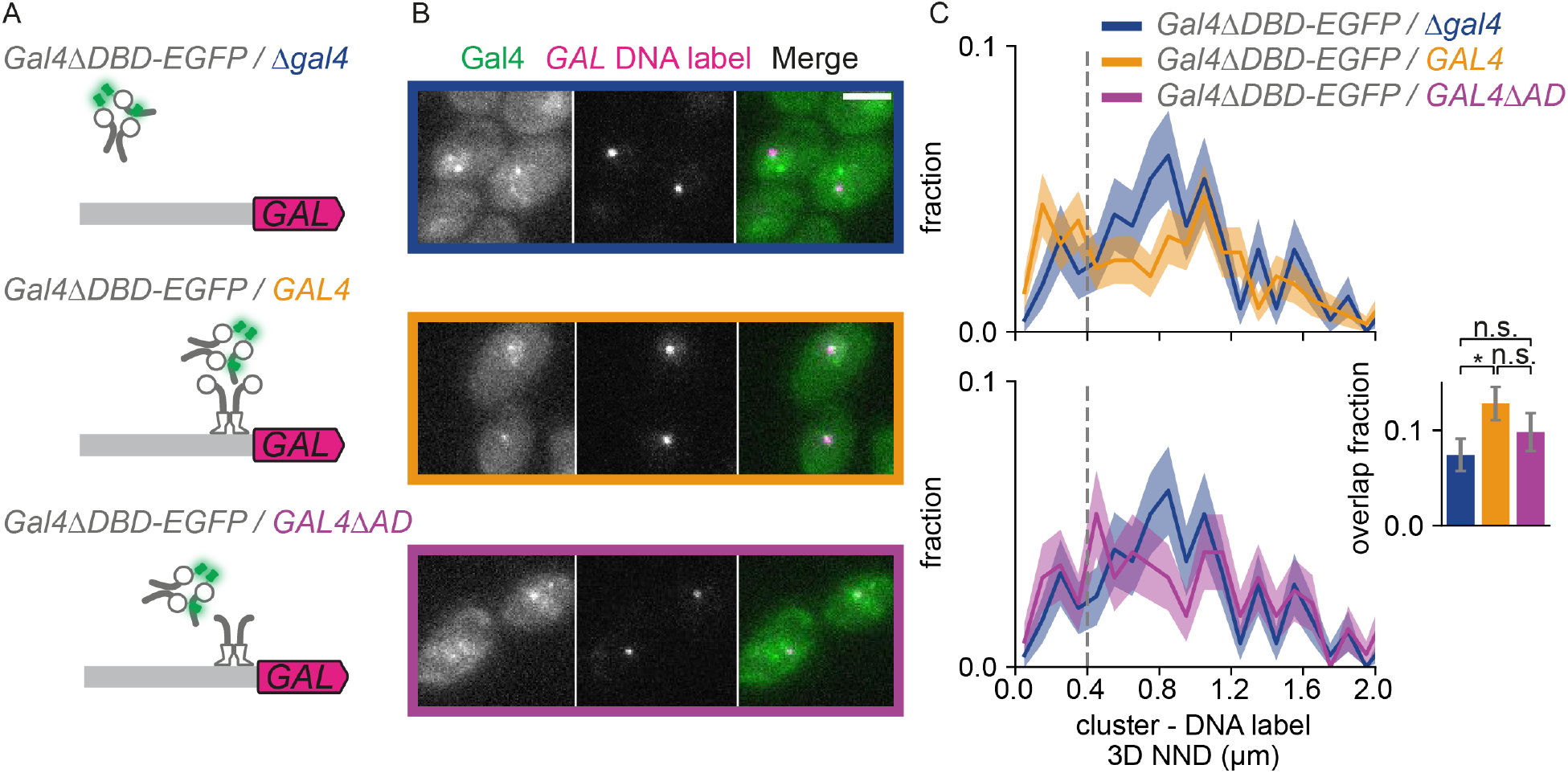
Gal4 self-interactions are sufficient to recruit Gal4 to target genes. **A.** Schematic representation of the *GAL* locus with DNA label (magenta) and diploid yeast strains expressing Gal4ΔDBD-EGFP from one allele and on the other allele expressing either *gal4*Δ (*BPSV40-GAL4(Δ2-94)-EGFP/gal4Δ*, blue), WT Gal4 (*BPSV40-GAL4(Δ1-94)-EGFP/GAL4*, orange), or Gal4ΔAD (*BPSV40-GAL4(Δ1-94)-EGFP/GAL4(Δ768-881)*, purple). **B.** Representative images of dual-color fluorescence imaging to determine the colocalization between Gal4ΔDBD-EGFP clusters (green) and the *GAL* DNA label (magenta) in the same yeast strains as **A.** after 30 minutes of induction (galactose and raffinose). Images are a single *z*-slice of a representative group of cells. **C.** Top: distribution of 3D nearest neighbor distances (NND) between the *GAL* DNA label and the closest Gal4ΔDBD-EGFP cluster for *GAL4ΔDBD-EGFP/gal4Δ* (blue, 300 cells) and *GAL4*Δ*DBD-EGFP/GAL4* (orange, 423 cells). Bottom: same as top, for *Gal4*Δ*DBD-EGFP/gal4*Δ (blue, same dataset as top panel) and *GAL4*Δ*DBD-EGFP/GAL4*Δ*AD* (purple, 256 cells). Shaded regions represent SEM based on 1000 bootstrap repeats. Vertical dashed line indicates 400 nm threshold used to discriminate between overlapping and non-overlapping clusters. Inset shows fraction of DNA-label containing cells with an overlapping cluster. Error bars represent SEM based on 1000 bootstrap repeats. Significance determined by Fisher’s exact test; n.s.: not significant; *: p < 0.05.

To independently validate these findings, we labeled the *GAL10* RNA with 14 repeats of PP7 hairpins to visualize the *GAL10* transcription site (TS). When the PP7 hairpin sequences at the 5’ of *GAL10* are transcribed, the nascent RNA is specifically bound by the PP7 coat protein, fused to a fluorescent protein ^60,61^. The *GAL10* TS is visible in the microscopy images as a bright nuclear spot. To measure recruitment of Gal4 to the active TS, we expressed either WT Gal4-EGFP or Gal4ΔDBD-EGFP from one allele and additional unlabeled Gal4 from a second allele to ensure transcriptional activity. For WT Gal4, around 75 ± 4% of *GAL10* TSs overlapped with a Gal4 cluster (Figure S6C-D). This overlap is higher than the overlap with the *GAL* DNA label (60 ± 4%), which may be partly explained by the fact that the RNA label is inside the locus rather than adjacent to it, resulting in smaller NNDs. Additionally, the higher overlap could hint at enrichment of colocalized Gal4 clusters when *GAL10* is active. Gal4 clusters that overlapped with the *GAL10* TS were brighter than Gal4 clusters that did not overlap (Figure S6F), in agreement with our findings using the *GAL* DNA label (Figure S2H). For Gal4ΔDBD-EGFP, a clear peak of close-proximity clusters was observed in the NND distribution, and approximately 43 ± 4% of *GAL10* TSs overlapped with a Gal4ΔDBD-EGFP cluster. These results are in line with our model that additional non-DNA-bound Gal4 molecules are recruited to target loci via protein-protein interactions, and that this recruitment is dependent on the Gal4 CR and/or AD. Transient self-interactions underlying Gal4 clusters are thus able to recruit Gal4 to its target genes.

### Colocalization of an active gene with a Gal4 cluster does not change transcriptional output

We next sought to understand how Gal4 clustering affects transcription activation. To link Gal4 clustering to the transcriptional output of its target genes, we used the PP7-*GAL10* RNA labeled strain to quantify transcription of the endogenous *GAL10* gene in living cells. In the microscopy images, the intensity of the TS is a measure for the number of nascent RNAs. To understand whether the colocalization of Gal4 is correlated to transcription levels, we compared the intensity of the *GAL10* TS in cells with and without an overlapping Gal4 cluster. This analysis showed that *GAL10* TSs that overlapped with a Gal4 cluster had the same intensity as non-overlapping TSs (Figure S6E). The presence of a Gal4 cluster at a target gene is thus uncorrelated with the transcription level of the target gene.

### Gal4 self-interactions are insufficient to activate transcription

The finding that Gal4 clustering does not correlate with the transcriptional activity of a target gene (Figures 1, 2, S6E) suggests that the Gal4 molecules in a cluster may not contribute to transcription activation. To test this further, we made use of the Gal4ΔAD and Gal4ΔDBD mutants that individually are unable to activate transcription as shown by their inability to grow on galactose-containing plates (Figure 6A). It is well-known that, when brought in close proximity, the Gal4 DBD and AD can function as transcriptional activator, even when they are part of different fusion proteins. This mechanism has been exploited in the classical Yeast-Two-Hybrid system to detect protein-protein interactions^62^. Our experiments showed that the Gal4ΔAD is able to recruit the Gal4ΔDBD to the Gal4 locus (Figure 5). We reasoned that if clustered Gal4ΔDBD molecules contribute to transcription activation, transient interactions between the DBD of the Gal4ΔAD mutant and the AD of the Gal4ΔDBD mutant within a cluster should rescue their transcriptional inactivity. However, coexpression of Gal4ΔAD and Gal4ΔDBD in a diploid yeast strain did not rescue the ability to activate *GAL* gene expression, as evidenced by their inability to grow on galactose-containing plates (Figure 6A). Despite being localized at the *GAL* genes, the AD of the Gal4ΔDBD mutant was unable to activate transcription (Figure 6B), suggesting that transient self-interactions are insufficient to activate gene expression. Moreover, the lack of rescue suggests that non-DNA-bound molecules in a cluster do not activate transcription.

**Figure 6:**
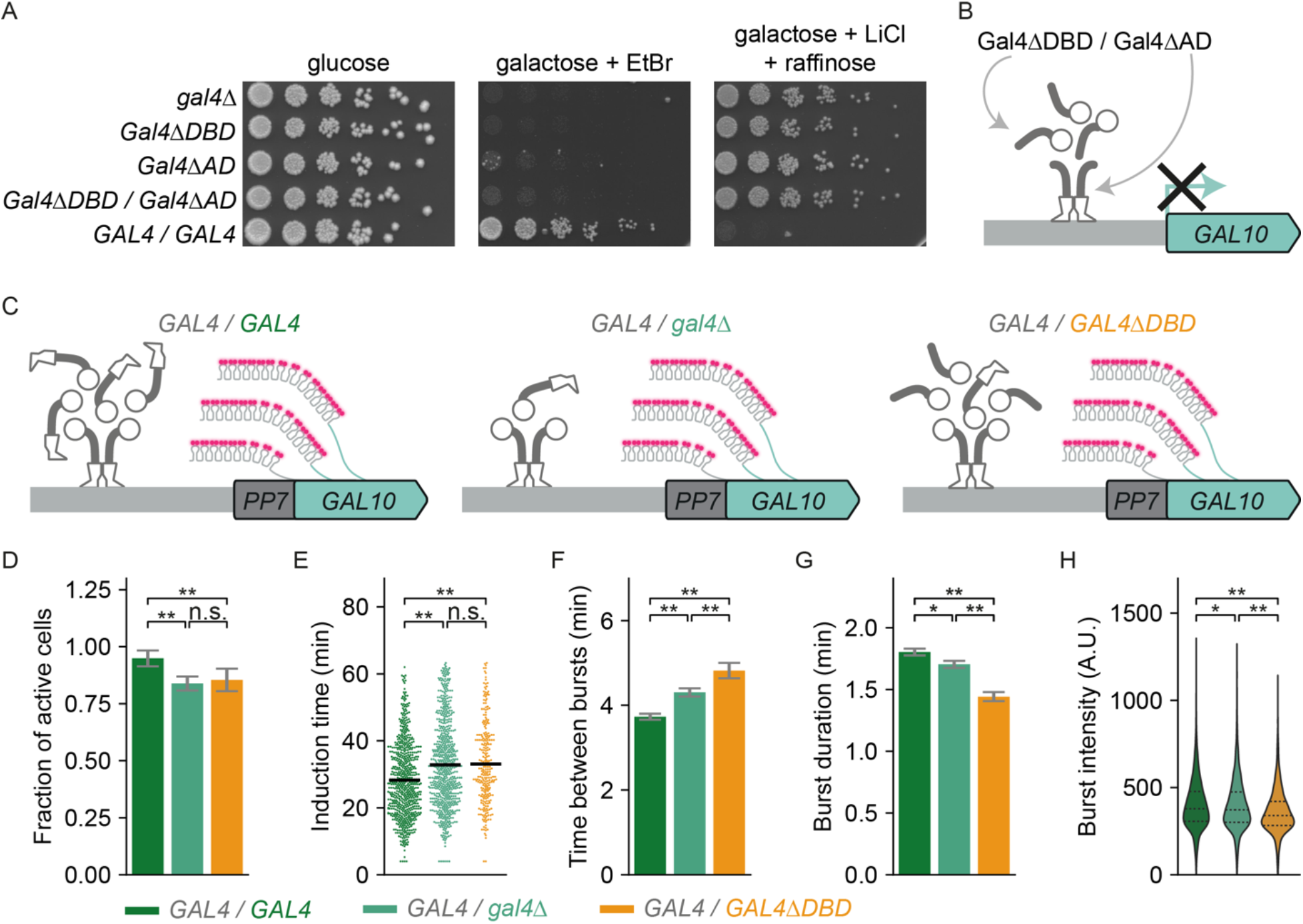
Gal4 self-interactions are insufficient to activate *GAL* gene transcription. **A.** Growth assay of indicated Gal4-EGFP mutants to assess their galactose metabolism capability. Shown are 5-fold serial dilutions on YEP + 2% glucose (dilution control), YEP + 2% galactose + 20 μg/mL ethidium bromide (growth = functional galactose metabolism) and YEP + 2% raffinose + 2% galactose + 40 mM lithium chloride + 0.003% methionine (no growth = functional galactose metabolism). **B.** Schematic representation of impaired *GAL* gene transcription in a diploid yeast strain expressing Gal4ΔAD from one allele and Gal4ΔDBD from the other allele (*GAL4(Δ768-881)-EGFP/BPSV40-GAL4(Δ2-94)-EGFP*). **C.** Schematic representation of the transcribed *PP7*-*GAL10* gene in diploid yeast strains expressing WT Gal4 from one allele and on the other allele expressing either WT Gal4 (*GAL4/GAL4,* green), Gal4Δ (*GAL4/gal4Δ*, light green) or Gal4ΔDBD (*GAL4/GAL4ΔDBD*, orange). Nascent *GAL10* transcripts are fluorescently labeled by binding of the PP7-coat protein-ymScarletI to the PP7-stem loops (magenta). **D-H.** Quantification of *PP7-GAL10* transcription in yeast strains described in **C.** with a total of 783, 889 and 350 cells, respectively. Cells were imaged during one hour of induction with galactose after growth in uninduced conditions (raffinose). **D.** Fraction of cells that activate *PP7*-*GAL10* transcription within one hour after induction. Error bars are propagated statistical errors in the number of active and inactive cells. Significance determined by Fisher’s exact test; n.s.: not significant; **: p < 0.01. **E.** Distributions of induction times of *PP7*-*GAL10* transcription. Black horizontal line: average as determined by 1000 bootstrap repeats. **F.** Average time between consecutive bursts and **G.** average burst duration of *PP7*-*GAL10* transcription, in which average and errors are determined by 1000 bootstrap repeats. **H.** Distribution of burst intensities of *PP7*-*GAL10* transcription. Black dashed lines show each quartile of the distribution. For **F-H** only cells with active *PP7-GAL10* transcription were included (673, 669 and 265 cells, respectively). Significance is determined by bootstrap hypothesis testing^79^; n.s.: not significant; *: p < 0.05; **: p < 0.01.

### Gal4 self-interactions at target genes can inhibit transcription

To further dissect the function of clustered non-DNA-bound Gal4 molecules in gene activation, we asked if clustered TF molecules that cannot bind DNA can enhance transcription activation in the presence of WT Gal4. We compared *GAL10* transcription between three diploid strains, all expressing WT Gal4 to activate transcription, and on the other allele either *GAL4*, *gal4*Δ or *GAL4*Δ*DBD* (Figure 6C). To quantify transcriptional activity, *GAL10* transcription was measured in live cells by PP7-*GAL10* imaging directly upon galactose addition. Compared to the cells with two *GAL4* gene copies (*GAL4/GAL4*), cells with one *GAL4* gene copy (*GAL4*/*gal4*Δ) had a slightly lower active fraction and took a longer time to activate *GAL10* after induction with galactose. Additionally, transcriptional bursts were shorter and time between consecutive bursts was longer (Figure 6D-H, S6G-I). If clustering of non-DNA-bound Gal4 molecules at a target site contributes to target gene transcription, nascent *GAL10* transcription in cells expressing Gal4Δ*DBD* on the second allele (*GAL4*/*GAL4*Δ*DBD*) should be increased compared to cells with only one Gal4 copy (*GAL4*/*gal4*Δ). However, live-cell imaging revealed that the presence of the Gal4ΔDBD mutant decreased the transcriptional output of *GAL10* even further, as we observed even shorter transcriptional bursts and even longer time between consecutive bursts, as well as a decrease in the burst intensity (Figure 6D-H, S6G-I). Therefore, we conclude that although the Gal4ΔDBD molecules can be recruited to a target site through self-interactions with WT Gal4 (Figure 5), these molecules do not contribute to transcription activation, and even inhibit gene transcription (Figure 6). Overall, our results indicate that the self-interactions that mediate TF clustering facilitate TF localization to target genes, but may also negatively influence transcription by inhibiting gene activation.

## DISCUSSION

In this manuscript we used quantitative imaging in living cells to understand how clustering of the paradigm TF Gal4 is regulated and how it influences transcription factor function. We found that Gal4 cluster abundance, size and intensity are regulated in different growth conditions through at least two mechanisms: (1) Glucose-repression of Gal4 expression levels decreased cluster abundance and size, and (2) Gal80-mediated Gal4 inhibition decreased Gal4 cluster abundance, size and density. In contrast, Gal4 clustering is largely unaffected by the well-established interactions with the Mediator complex through the subunit Med15. Moreover, DNA binding enhances cluster formation, and enables clusters to become denser and smaller, especially at the multiple Gal4 UASs of the *GAL1-GAL10-GAL7* locus. Gal4 clustering is also facilitated by, but not completely dependent on the CR and AD, which contain several IDRs. Through combinations of various truncation mutants, we showed that clusters consist of DNA-bound Gal4 molecules as well as non-DNA bound Gal4 molecules that are likely recruited to target genes through proteinprotein interactions of the CR and ADs. However, once at the target site, these non-DNA bound TF molecules do not contribute to gene activation, and may in fact inhibit gene transcription.

### Gal4 cluster formation

TF clusters have previously been described to be driven by both homotypic and heterotypic interactions^63^. For Gal4, several findings suggest that clusters are formed mostly by homotypic rather than heterotypic interactions: (1) Clustering was not reduced by loss of interactions with Med15. (2) Removal of the miniAD presumably results in loss of most heterotypic interactions with the transcriptional machinery^64^, but yielded similar clustering as WT. (3) Gal4 without a DBD can be recruited to the *GAL* locus by WT Gal4 or Gal4 without an AD. To show the contribution of homotypic self-interactions or heterotypic interactions of TFs with other factors, previous studies have used *in vitro* analysis with purified proteins^4,11,16^. Gal4, however, has been notoriously difficult to purify as a full-length protein, preventing us from performing such *in vitro* experiments^65^. Regardless, our imaging tools create the possibility to test in the future how Gal4 clustering is regulated by other Gal4 interactors *in vivo*, such as SAGA, TBP or TFIIB^64,66,67^. Moreover, it will be interesting to explore whether such factors co-cluster with Gal4 near target genes, as was observed for Hsf1 condensates^29^. Although the role of heterotypic interactions in Gal4 remains unclear, our data suggests that Gal4 homotypic self-interactions are an important contributor to Gal4 clustering.

Previously, homo- and heterotypic multivalent interactions have been linked to the formation of liquid-liquid phase separated condensates^4,63^. The experiments with increasing glucose concentration show that increasing the Gal4 concentration increases the cluster abundance and size, but has minor effects on the cluster density. These results are consistent with the simplest liquid-liquid phase separation model, which postulates that at higher concentrations, more clusters are formed and that clusters merge into larger clusters with the same concentration inside the cluster^68,69^. Although our concentration-gradient is consistent with this model, further tests would be required to understand the role of liquid-liquid phase separation in Gal4 cluster formation.

### Gal4 cluster regulation by Gal80

In inactive conditions, Gal4 clustering is inhibited by its repressor Gal80. Gal80 binds to the Gal4 AD, resulting in a structured conformation of the AD and masking it for interactions with transcription cofactors such as Mediator^70,71^. Loss of Med15 did not majorly affect Gal4 clustering, indicating that the inhibitory effect of Gal80 on clustering is likely mediated by preventing homotypic interactions rather than by preventing Mediator interactions. Gal80 prevents these homotypic interactions likely by shielding a larger region than the AD, as Gal4 clusters also formed efficiently in the Gal4ΔAD mutant.

A previous study showed that Gal80 also localizes in nuclear clusters that dissociate upon galactose addition^72^. Self-association of Gal80 was proposed to be required for Gal4 repression by Gal80, explaining why genes with multiple Gal4 UASs may be more efficiently repressed than genes with a single Gal4 UAS^73^. The corollary of this model is that the inhibitory capacity of Gal80 may be enhanced by Gal4 clustering, but that Gal80 simultaneously inhibits Gal4 clustering, thereby limiting its own inhibitory capacity. Conversely, if Gal80 and Gal4 oligomerization enhances their function, perhaps the formation of clusters is merely a consequence of this function.

### Role of DNA binding and IDRs in Gal4 clustering

Clustering is often suggested to be related to IDR-mediated protein-protein interactions^4,12,17^. However, the exact role of different TF domains remains unclear and may differ per TF. For example, for the pioneer factor Sox2 clustering depends on the DNA binding domain but not the activation domain^18^, suggesting that clustering is the result of the spatial proximity of multiple TF binding sites. On the other hand, clustering of the Oct4 TF depends on the IDRs in the ADs^4^. Here, we find that for Gal4, clustering depends both on DNA binding and the IDRs.

When the Gal4 DBD is deleted or mutated, clusters are still observed, although they are much less abundant and less dense (Figure 3). These effects on cluster abundance are larger compared to the UASscr mutant, indicating that DNA binding at single sites considerably contributes to cluster formation. However, the presence of multiple Gal4 UASs in individual promoters increases the cluster density. DNA binding may form a scaffold that lowers the propensity for cluster formation, which may be enhanced if multiple UASs are placed adjacently, similar to enhancers in mammalian cells^17,74^.

Besides DNA binding, Gal4 clustering is also facilitated by the IDRs in the CR and AD. Removal of all predicted IDRs, leaving only the structured DBD and dimerization domain of Gal4 (Gal4-DBD-only), resulted in a large decrease in cluster density and abundance (Figure 4). However, some clusters are still observed upon IDR loss, suggesting that the structured DBD and/or dimerization domain contribute to clustering. In accordance, clustering of IDRs was reported to be driven by their multivalency rather than by their disorderedness^75^ and such multivalency may also be present in structured domains. Given that the dimerization domain is capable of interacting with a specific Med15 mutant (Gal11p)^49^, we speculate that the dimerization domain may be able to form homotypic and heterotypic interactions that promote clustering. Along these same lines, besides the predicted IDRs, the CR also contains a predicted structured region (Figure S5A) which may contribute to the clustering potential of the CR. We conclude that both DNA binding and IDRs contribute to Gal4 clustering.

### Self-interactions facilitate target search

In eukaryotic cells, TFs face a major challenge in finding their targets in millions of non-specific sequences. Our results indicate that self-interactions may facilitate this target search, in line with previous findings for several yeast and mammalian transcriptional regulators^24–28^. Although localization of Gal4 clusters is dependent on the sequence-specific Gal4 DBD, self-interactions between regions outside the DBD promote the formation and localization of Gal4 clusters to the *GAL* locus. We envision that once Gal4 molecules are bound to the DNA, their exposed IDRs allow for interaction with additional unbound Gal4 molecules, thereby creating a larger effective target size. Such a mechanism may ensure that the DBD of each Gal4 molecule does not need to probe all DNA sequences to find its binding site, and reduces the search space to areas where other Gal4 molecules have already found their target^13,76^.

In this model, clustering also creates cooperativity, as binding of subsequent molecules is enhanced after the first one is bound. This cooperative binding through clustering has been proposed to be important for mammalian super enhancers that contain many TF binding sites^22^. For Gal4, four target genes contain multiple binding sites, which include the genes encoding galactose-metabolic enzymes^47^. At these genes, cooperative binding from clustering may enable fast transcriptional response upon exposure to galactose and may ensure high Gal4 promoter occupancy and high expression. Conversely, several other Gal4 target promoters only contain one UAS and it remains to be established whether clustering and self-interactions are beneficial for target search for these genes. Since a cluster located near a promoter locally increases the concentration of Gal4 molecules, we can imagine that, once a Gal4 molecule is released from the DNA, quick rebinding of another Gal4 from the nearby cluster to the DNA may increase the overall time that the promoter is occupied. In addition, self-interactions between the cluster and the bound Gal4 molecules could increase the dwell time of the bound molecule^12,21^. It will be interesting in the future to examine how self-interactions and clustering influence Gal4 binding kinetics at different genes and at the single-molecule level.

### Clustered Gal4 molecules do not activate transcription

Although Gal4 clusters overlap with transcriptionally active loci, *GAL* gene transcription is not supported by additional non-DNA-bound Gal4 molecules in the cluster. In a yeast-two-hybrid-like setup, DNA-bound Gal4ΔAD without an AD that recruited a Gal4ΔDBD without an DBD to the target gene could not activate transcription. Likely, the classical yeast-two-hybrid setup to detect protein-protein interaction only detects stable lock-and-key interactions and not “fuzzy” interactions such as those facilitated by TF clustering. Moreover, the presence of extra non-DNA-bound mutant Gal4 molecules at the target gene inhibited rather than enhanced transcription activation. Based on these findings, we conclude Gal4 needs to be DNA-bound to activate transcription.

Our experiments with truncation mutants allowed us to establish that non-DNA bound molecules do not contribute to gene activation. We observe that these non-DNA bound molecules may inhibit transcription, but this could be due to a dominant-negative effect of the Gal4ΔDBD mutant. In these experiments, the concentration of DBDs within clusters is reduced, which may lower the on-rate compared to WT. Since such truncated proteins are not present in WT cells, we cannot directly test whether these results apply to WT clusters. However, given that transcription inhibition from clustering has been observed previously^33,34^, it is conceivable that Gal4 clustering also inhibits gene expression in a WT context.

In this latter case, we speculate that the observed transcription inhibition inside clusters occurs through a titration mechanism called squelching^77^. Gal4 overexpression has been described to inhibit transcription by titrating the transcriptional machinery^77^. The high Gal4 concentration inside the cluster may cause competition between non-DNA-bound Gal4 with DNA-bound Gal4 for interactions with cofactors and the preinitiation complex. A similar transcription inhibition has recently been observed for the FUS-EWS fusion, where additional expression of the EWS domain decreased the ability of the endogenous FUS-EWS fusion to active transcription^33^. Here, additional non-DNA-bound EWS may squelch DNA bound FUS-EWS. In addition, our model is in line with the recent finding that chemically-induced clustering caused a negative correlation between cluster intensity and transcriptional output^32^. These findings suggests that self-interactions that cause clustering may compete with, rather than facilitate, recruitment of the transcriptional machinery.

### Clustering acts as a double-edged sword

Transcription factors need to perform two major steps during transcription activation: binding to their target genes and activating transcription by recruiting the transcriptional machinery. Our results show that Gal4 clustering enables DNA binding but inhibits transcription activation. This finding suggests that cells need to balance these positive and negative aspects of clustering for proper gene expression. Depending on the propensity to engage in homotypic and heterotypic interactions, this balance may shift for different transcription factors. Such a balance may perhaps also clarify why clustering and liquid-liquid phase separation may be beneficial in certain systems and inhibitory in others. Our quantitative imaging approach will open up new avenues to explore this balance for other TFs in the future.

### Limitations of the study

A current limitation of this study is that we could only reliably analyze clusters of Gal4 mutants that showed relatively low expression. In several mutants we tested, the expression level increased to a level that made it challenging to distinguish the background from the many clusters in the small yeast nucleus. We did not include these mutants in this manuscript. Our attempts to perform super resolution imaging of Gal4 clusters to provide insight into their size and structure failed because our Gal4 cluster detection was limited to live cells. Gal4 clusters were not visible in fixed samples using various fixation chemicals, suggesting that Gal4 clusters are too dynamic to be not efficiently crosslinked^78^. We also note that Gal4 cluster detection required high excitation power and HILO illumination, perhaps explaining why previous microscopy studies were not successful in detecting Gal4 clusters^57,72^. These high excitation powers quickly resulted in bleaching, precluding us from studying Gal4 clusters for prolonged times. Future improvements of fluorescent proteins and microscopy systems may allow for longer imaging to determine the dynamics of Gal4 clusters.

## Acknowledgements

We thank the Amir Aharoni laboratory for plasmids and the Fred van Leeuwen laboratory for western blot antibodies. We thank the Research High Performance Computing Facility of the NKI for assistance. We thank members of the T.L.L. and Fred van Leeuwen laboratories for helpful discussions, Bas van Steensel for guidance and Elzo de Wit for critical reading of the manuscript. This work was supported by the Netherlands Organization for Scientific Research (NWO, 016.Veni.192.071 (I.B.) and gravitation program CancerGenomiCs.nl (T.L.L.)), Oncode Institute (T.L.L.), which is partly financed by the Dutch Cancer Society, and the European Research Council (ERC Starting Grant 755695 BURSTREG (T.L.L.)) and an institutional grant of the Dutch Cancer Society and of the Dutch Ministry of Health, Welfare and Sport.

## Author contributions

Conceptualization, J.V.W.M., H.P.P., W.J.J. and T.L.L.; Methodology, J.V.W.M., I.B., W.J.J., H.P.P. and T.L.L.; Software, J.V.W.M., W.P. and T.L.L.; Formal Analysis, J.V.W.M. and I.B.; Investigation, J.V.W.M., I.B., W.J.J. and H.P.P.; Data Curation, J.V.W.M. and W.P.P.; Writing – Original Draft, J.V.W.M., I.B. and T.L.L.; Writing – Review & Editing, J.V.W.M, W.P., I.B., W.J.J., H.P.P. and T.L.L; Visualization, J.V.W.M., I.B., W.J.J. and T.L.L.; Supervision, T.L.L.; Funding Acquisition, I.B. and T.L.L.

## REFERENCES

1. Ptashne, M., and Gann, A. (1997). Transcriptional activation by recruitment. Nature 386, 569–577. 10.1038/386569a0.

2. van Steensel, B., Brink, M., van der Meulen, K., van Binnendijk, E.P., Wansink, D.G., de Jong, L., de Kloet, E.R., and van Driel, R. (1995). Localization of the glucocorticoid receptor in discrete clusters in the cell nucleus. J. Cell Sci. 108 (Pt 9), 3003–3011. 10.1242/jcs.108.9.3003.

3. Boehning, M., Dugast-Darzacq, C., Rankovic, M., Hansen, A.S., Yu, T., Marie-Nelly, H., McSwiggen, D.T., Kokic, G., Dailey, G.M., Cramer, P., et al. (2018). RNA polymerase II clustering through carboxy-terminal domain phase separation. Nat. Struct. Mol. Biol. 25, 833–840. 10.1038/s41594-018-0112-y.

4. Boija, A., Klein, I.A., Sabari, B.R., Dall’Agnese, A., Coffey, E.L., Zamudio, A.V., Li, C.H., Shrinivas, K., Manteiga, J.C., Hannett, N.M., et al. (2018). Transcription Factors Activate Genes through the Phase-Separation Capacity of Their Activation Domains. Cell 175, 1842–1855.e16. 10.1016/j.cell.2018.10.042.

5. Cho, W.-K., Spille, J.-H., Hecht, M., Lee, C., Li, C., Grube, V., and Cisse, I.I. (2018). Mediator and RNA polymerase II clusters associate in transcription-dependent condensates. Science 361, 412–415. 10.1126/science.aar4199.

6. Dufourt, J., Trullo, A., Hunter, J., Fernandez, C., Lazaro, J., Dejean, M., Morales, L., Nait-Amer, S., Schulz, K.N., Harrison, M.M., et al. (2018). Temporal control of gene expression by the pioneer factor Zelda through transient interactions in hubs. Nat. Commun. 9, 5194. 10.1038/s41467-018-07613-z.

7. Li, J., Dong, A., Saydaminova, K., Chang, H., Wang, G., Ochiai, H., Yamamoto, T., and Pertsinidis, A. (2019). Single-Molecule Nanoscopy Elucidates RNA Polymerase II Transcription at Single Genes in Live Cells. Cell 178, 491–506.e28. 10.1016/j.cell.2019.05.029.

8. Liu, Z., Legant, W.R., Chen, B.-C., Li, L., Grimm, J.B., Lavis, L.D., Betzig, E., and Tjian, R. (2014). 3D imaging of Sox2 enhancer clusters in embryonic stem cells. eLife 3, e04236. 10.7554/eLife.04236.

9. Mir, M., Reimer, A., Haines, J.E., Li, X.-Y., Stadler, M., Garcia, H., Eisen, M.B., and Darzacq, X. (2017). Dense Bicoid hubs accentuate binding along the morphogen gradient. Genes Dev. 31, 1784–1794. 10.1101/gad.305078.117.

10. Mir, M., Stadler, M.R., Ortiz, S.A., Hannon, C.E., Harrison, M.M., Darzacq, X., and Eisen, M.B. (2018). Dynamic multifactor hubs interact transiently with sites of active transcription in Drosophila embryos. eLife 7, e40497. 10.7554/eLife.40497.

11. Sabari, B.R., Dall’Agnese, A., Boija, A., Klein, I.A., Coffey, E.L., Shrinivas, K., Abraham, B.J., Hannett, N.M., Zamudio, A.V., Manteiga, J.C., et al. (2018). Coactivator condensation at super-enhancers links phase separation and gene control. Science 361, eaar3958. 10.1126/science.aar3958.

12. Chong, S., Dugast-Darzacq, C., Liu, Z., Dong, P., Dailey, G.M., Cattoglio, C., Heckert, A., Banala, S., Lavis, L., Darzacq, X., et al. (2018). Imaging dynamic and selective low-complexity domain interactions that control gene transcription. Science 361, eaar2555. 10.1126/science.aar2555.

13. Ferrie, J.J., Karr, J.P., Tjian, R., and Darzacq, X. (2022). “Structure”-function relationships in eukaryotic transcription factors: The role of intrinsically disordered regions in gene regulation. Mol. Cell 82, 3970–3984. 10.1016/j.molcel.2022.09.021.

14. Kwon, I., Kato, M., Xiang, S., Wu, L., Theodoropoulos, P., Mirzaei, H., Han, T., Xie, S., Corden, J.L., and McKnight, S.L. (2013). Phosphorylation-Regulated Binding of RNA Polymerase II to Fibrous Polymers of Low-Complexity Domains. Cell 155, 1049–1060. 10.1016/j.cell.2013.10.033.

15. Tuttle, L.M., Pacheco, D., Warfield, L., Wilburn, D.B., Hahn, S., and Klevit, R.E. (2021). Mediator subunit Med15 dictates the conserved “fuzzy” binding mechanism of yeast transcription activators Gal4 and Gcn4. Nat. Commun. 12, 2220. 10.1038/s41467-021-22441-4.

16. Wang, W., Qiao, S., Li, G., Cheng, J., Yang, C., Zhong, C., Stovall, D.B., Shi, J., Teng, C., Li, D., et al. (2022). A histidine cluster determines YY1-compartmentalized coactivators and chromatin elements in phase-separated enhancer clusters. Nucleic Acids Res. 50, 4917–4937. 10.1093/nar/gkac233.

17. Shrinivas, K., Sabari, B.R., Coffey, E.L., Klein, I.A., Boija, A., Zamudio, A.V., Schuijers, J., Hannett, N.M., Sharp, P.A., Young, R.A., et al. (2019). Enhancer Features that Drive Formation of Transcriptional Condensates. Mol. Cell 75, 549–561.e7. 10.1016/j.molcel.2019.07.009.

18. Li, J., Hsu, A., Hua, Y., Wang, G., Cheng, L., Ochiai, H., Yamamoto, T., and Pertsinidis, A. (2020). Single-gene imaging links genome topology, promoter-enhancer communication and transcription control. Nat. Struct. Mol. Biol. 27, 1032–1040. 10.1038/s41594-020-0493-6.

19. Stortz, M., Pecci, A., Presman, D.M., and Levi, V. (2020). Unraveling the molecular interactions involved in phase separation of glucocorticoid receptor. BMC Biol. 18, 59. 10.1186/s12915-020-00788-2.

20. Ma, L., Gao, Z., Wu, J., Zhong, B., Xie, Y., Huang, W., and Lin, Y. (2021). Co-condensation between transcription factor and coactivator p300 modulates transcriptional bursting kinetics. Mol. Cell 81, 1682–1697.e7. 10.1016/j.molcel.2021.01.031.

21. de Jonge, W.J., Patel, H.P., Meeussen, J.V.W., and Lenstra, T.L. (2022). Following the tracks: How transcription factor binding dynamics control transcription. Biophys. J. 121, 1583–1592. 10.1016/j.bpj.2022.03.026.

22. Hnisz, D., Shrinivas, K., Young, R.A., Chakraborty, A.K., and Sharp, P.A. (2017). A Phase Separation Model for Transcriptional Control. Cell 169, 13–23. 10.1016/j.cell.2017.02.007.

23. Mazzocca, M., Fillot, T., Loffreda, A., Gnani, D., and Mazza, D. (2021). The needle and the haystack: single molecule tracking to probe the transcription factor search in eukaryotes. Biochem. Soc. Trans. 49, 1121–1132. 10.1042/BST20200709.

24. Brodsky, S., Jana, T., Mittelman, K., Chapal, M., Kumar, D.K., Carmi, M., and Barkai, N. (2020). Intrinsically Disordered Regions Direct Transcription Factor In Vivo Binding Specificity. Mol. Cell 79, 459–471.e4. 10.1016/j.molcel.2020.05.032.

25. Chen, Y., Cattoglio, C., Dailey, G.M., Zhu, Q., Tjian, R., and Darzacq, X. (2022). Mechanisms governing target search and binding dynamics of hypoxia-inducible factors. eLife 11, e75064. 10.7554/eLife.75064.

26. Hansen, A.S., Amitai, A., Cattoglio, C., Tjian, R., and Darzacq, X. (2020). Guided nuclear exploration increases CTCF target search efficiency. Nat. Chem. Biol. 16, 257–266. 10.1038/s41589-019-0422-3.

27. Kent, S., Brown, K., Yang, C.-H., Alsaihati, N., Tian, C., Wang, H., and Ren, X. (2020). Phase-Separated Transcriptional Condensates Accelerate Target-Search Process Revealed by Live-Cell Single-Molecule Imaging. Cell Rep. 33, 108248. 10.1016/j.celrep.2020.108248.

28. Wollman, A.J., Shashkova, S., Hedlund, E.G., Friemann, R., Hohmann, S., and Leake, M.C. (2017). Transcription factor clusters regulate genes in eukaryotic cells. eLife 6, e27451. 10.7554/eLife.27451.

29. Chowdhary, S., Kainth, A.S., Paracha, S., Gross, D.S., and Pincus, D. (2022). Inducible transcriptional condensates drive 3D genome reorganization in the heat shock response. Mol. Cell, S1097-2765(22)00970-4. 10.1016/j.molcel.2022.10.013.

30. Wei, M.-T., Chang, Y.-C., Shimobayashi, S.F., Shin, Y., Strom, A.R., and Brangwynne, C.P. (2020). Nucleated transcriptional condensates amplify gene expression. Nat. Cell Biol. 22, 1187–1196. 10.1038/s41556-020-00578-6.

31. Schneider, N., Wieland, F.-G., Kong, D., Fischer, A.A.M., Hörner, M., Timmer, J., Ye, H., and Weber, W. (2021). Liquid-liquid phase separation of light-inducible transcription factors increases transcription activation in mammalian cells and mice. Sci. Adv. 7, eabd3568. 10.1126/sciadv.abd3568.

32. Wu, J., Chen, B., Liu, Y., Ma, L., Huang, W., and Lin, Y. (2022). Modulating gene regulation function by chemically controlled transcription factor clustering. Nat. Commun. 13, 2663. 10.1038/s41467-022-30397-2.

33. Chong, S., Graham, T.G.W., Dugast-Darzacq, C., Dailey, G.M., Darzacq, X., and Tjian, R. (2022). Tuning levels of low-complexity domain interactions to modulate endogenous oncogenic transcription. Mol. Cell 82, 2084–2097.e5. 10.1016/j.molcel.2022.04.007.

34. Trojanowski, J., Frank, L., Rademacher, A., Mücke, N., Grigaitis, P., and Rippe, K. (2022). Transcription activation is enhanced by multivalent interactions independent of phase separation. Mol. Cell 82, 1878–1893.e10. 10.1016/j.molcel.2022.04.017.

35. Traven, A., Jelicic, B., and Sopta, M. (2006). Yeast Gal4: a transcriptional paradigm revisited. EMBO Rep. 7, 496–499. 10.1038/sj.embor.7400679.

36. Ma, J., and Ptashne, M. (1987). The carboxy-terminal 30 amino acids of GAL4 are recognized by GAL80. Cell 50, 137–142. 10.1016/0092-8674(87)90670-2.

37. Ricci-Tam, C., Ben-Zion, I., Wang, J., Palme, J., Li, A., Savir, Y., and Springer, M. (2021). Decoupling transcription factor expression and activity enables dimmer switch gene regulation. Science 372, 292–295. 10.1126/science.aba7582.

38. Sellick, C.A., Campbell, R.N., and Reece, R.J. (2008). Chapter 3 Galactose Metabolism in Yeast—Structure and Regulation of the Leloir Pathway Enzymes and the Genes Encoding Them. In International Review of Cell and Molecular Biology (Academic Press), pp. 111–150. 10.1016/S1937-6448(08)01003-4.

39. Tokunaga, M., Imamoto, N., and Sakata-Sogawa, K. (2008). Highly inclined thin illumination enables clear single-molecule imaging in cells. Nat. Methods 5, 159–161. 10.1038/nmeth1171.

40. Dovrat, D., Dahan, D., Sherman, S., Tsirkas, I., Elia, N., and Aharoni, A. (2018). A Live-Cell Imaging Approach for Measuring DNA Replication Rates. Cell Rep. 24, 252–258. 10.1016/j.celrep.2018.06.018.

41. Donovan, B.T., Huynh, A., Ball, D.A., Patel, H.P., Poirier, M.G., Larson, D.R., Ferguson, M.L., and Lenstra, T.L. (2019). Live-cell imaging reveals the interplay between transcription factors, nucleosomes, and bursting. EMBO J. 38. 10.15252/embj.2018100809.

42. Shi, Y., Chen, J., Zeng, W.-J., Li, M., Zhao, W., Zhang, X.-D., and Yao, J. (2021). Formation of nuclear condensates by the Mediator complex subunit Med15 in mammalian cells. BMC Biol. 19, 245. 10.1186/s12915-021-01178-y.

43. Griggs, D.W., and Johnston, M. (1991). Regulated expression of the GAL4 activator gene in yeast provides a sensitive genetic switch for glucose repression. Proc. Natl. Acad. Sci. U. S. A. 88, 8597–8601. 10.1073/pnas.88.19.8597.

44. Laughon, A., and Gesteland, R.F. (1982). Isolation and preliminary characterization of the GAL4 gene, a positive regulator of transcription in yeast. Proc. Natl. Acad. Sci. U. S. A. 79, 6827–6831. 10.1073/pnas.79.22.6827.

45. Giniger, E., Varnum, S.M., and Ptashne, M. (1985). Specific DNA binding of GAL4, a positive regulatory protein of yeast. Cell 40, 767–774. 10.1016/0092-8674(85)90336-8.

46. Marmorstein, R., Carey, M., Ptashne, M., and Harrison, S.C. (1992). DNA recognition by GAL4: structure of a protein-DNA complex. Nature 356, 408–414. 10.1038/356408a0.

47. Rhee, H.S., and Pugh, B.F. (2011). Comprehensive Genome-wide Protein-DNA Interactions Detected at Single Nucleotide Resolution. Cell 147, 1408–1419. 10.1016/j.cell.2011.11.013.

48. Carey, M., Kakidani, H., Leatherwood, J., Mostashari, F., and Ptashne, M. (1989). An aminoterminal fragment of GAL4 binds DNA as a dimer. J. Mol. Biol. 209, 423–432. 10.1016/0022-2836(89)90007-7.

49. Hong, M., Fitzgerald, M.X., Harper, S., Luo, C., Speicher, D.W., and Marmorstein, R. (2008). Structural Basis for Dimerization in DNA Recognition by Gal4. Struct. Lond. Engl. 1993 16, 1019–1026. 10.1016/j.str.2008.03.015.

50. Silver, P.A., Keegan, L.P., and Ptashne, M. (1984). Amino Terminus of the Yeast GAL4 Gene Product is Sufficient for Nuclear Localization. Proc. Natl. Acad. Sci. U. S. A. 81, 5951–5955.

51. Silver, P.A., Chiang, A., and Sadler, I. (1988). Mutations that alter both localization and production of a yeast nuclear protein. Genes Dev. 2, 707–717. 10.1101/gad.2.6.707.

52. Hodel, A.E., Harreman, M.T., Pulliam, K.F., Harben, M.E., Holmes, J.S., Hodel, M.R., Berland, K.M., and Corbett, A.H. (2006). Nuclear Localization Signal Receptor Affinity Correlates with in Vivo Localization in Saccharomyces cerevisiae*. J. Biol. Chem. 281, 23545–23556. 10.1074/jbc.M601718200.

53. Jeličić, B., Nemet, J., Traven, A., and Sopta, M. (2014). e. FEMS Yeast Res. 14, 302–309. 10.1111/1567-1364.12106.

54. Burdach, J., Funnell, A.P.W., Mak, K.S., Artuz, C.M., Wienert, B., Lim, W.F., Tan, L.Y., Pearson, R.C.M., and Crossley, M. (2014). Regions outside the DNA-binding domain are critical for proper in vivo specificity of an archetypal zinc finger transcription factor. Nucleic Acids Res. 42, 276–289. 10.1093/nar/gkt895.

55. Lim, W.F., Burdach, J., Funnell, A.P.W., Pearson, R.C.M., Quinlan, K.G.R., and Crossley, M. (2016). Directing an artificial zinc finger protein to new targets by fusion to a non-DNA-binding domain. Nucleic Acids Res. 44, 3118–3130. 10.1093/nar/gkv1380.

56. Völkel, S., Stielow, B., Finkernagel, F., Stiewe, T., Nist, A., and Suske, G. (2015). Zinc Finger Independent Genome-Wide Binding of Sp2 Potentiates Recruitment of Histone-Fold Protein Nf-y Distinguishing It from Sp1 and Sp3. PLOS Genet. 11, e1005102. 10.1371/journal.pgen.1005102.

57. Mizutani, A., and Tanaka, M. (2003). Regions of GAL4 critical for binding to a promoter in vivo revealed by a visual DNA-binding analysis. EMBO J. 22, 2178–2187. 10.1093/emboj/cdg220.

58. Erdős, G., Pajkos, M., and Dosztányi, Z. (2021). IUPred3: prediction of protein disorder enhanced with unambiguous experimental annotation and visualization of evolutionary conservation. Nucleic Acids Res. 49, W297–W303. 10.1093/nar/gkab408.

59. Erdős, G., and Dosztányi, Z. (2020). Analyzing Protein Disorder with IUPred2A. Curr. Protoc. Bioinforma. 70, e99. 10.1002/cpbi.99.

60. Larson, D.R., Zenklusen, D., Wu, B., Chao, J.A., and Singer, R.H. (2011). Real-time observation of transcription initiation and elongation on an endogenous yeast gene. Science 332, 475–478. 10.1126/science.1202142.

61. Lenstra, T.L., and Larson, D.R. (2016). Single-Molecule mRNA Detection in Live Yeast. Curr. Protoc. Mol. Biol. 113, 14.24.1–14.24.15. 10.1002/0471142727.mb1424s113.

62. Fields, S., and Song, O. (1989). A novel genetic system to detect protein-protein interactions. Nature 340, 245–246. 10.1038/340245a0.

63. Mittag, T., and Pappu, R.V. (2022). A conceptual framework for understanding phase separation and addressing open questions and challenges. Mol. Cell 82, 2201–2214. 10.1016/j.molcel.2022.05.018.

64. Wu, Y., Reece, R.J., and Ptashne, M. (1996). Quantitation of putative activator-target affinities predicts transcriptional activating potentials. EMBO J. 15, 3951–3963.

65. Ding, W.V., and Johnston, S.A. (1997). The DNA binding and activation domains of Gal4p are sufficient for conveying its regulatory signals. Mol. Cell. Biol. 17, 2538–2549. 10.1128/MCB.17.5.2538.

66. Bhaumik, S.R., Raha, T., Aiello, D.P., and Green, M.R. (2004). In vivo target of a transcriptional activator revealed by fluorescence resonance energy transfer. Genes Dev. 18, 333–343. 10.1101/gad.1148404.

67. Melcher, K., and Johnston, S.A. (1995). GAL4 interacts with TATA-binding protein and coactivators. Mol. Cell. Biol. 15, 2839–2848. 10.1128/MCB.15.5.2839.

68. Alberti, S., Gladfelter, A., and Mittag, T. (2019). Considerations and challenges in studying liquid-liquid phase separation and biomolecular condensates. Cell 176, 419–434. 10.1016/j.cell.2018.12.035.

69. Shin, Y., Berry, J., Pannucci, N., Haataja, M.P., Toettcher, J.E., and Brangwynne, C.P. (2017). Spatiotemporal control of intracellular phase transitions using light-activated optoDroplets. Cell 168, 159–171.e14. 10.1016/j.cell.2016.11.054.

70. Kumar, P.R., Yu, Y., Sternglanz, R., Johnston, S.A., and Joshua-Tor, L. (2008). NADP regulates the yeast GAL induction system. Science 319, 1090–1092. 10.1126/science.1151903.

71. Thoden, J.B., Ryan, L.A., Reece, R.J., and Holden, H.M. (2008). The interaction between an acidic transcriptional activator and its inhibitor. The molecular basis of Gal4p recognition by Gal80p. J. Biol. Chem. 283, 30266–30272. 10.1074/jbc.M805200200.

72. Egriboz, O., Goswami, S., Tao, X., Dotts, K., Schaeffer, C., Pilauri, V., and Hopper, J.E. (2013). Self-Association of the Gal4 Inhibitor Protein Gal80 Is Impaired by Gal3: Evidence for a New Mechanism in the GAL Gene Switch. Mol. Cell. Biol. 33, 3667–3674. 10.1128/MCB.00646-12.

73. Bram, R.J., Lue, N.F., and Kornberg, R.D. (1986). A GAL family of upstream activating sequences in yeast: roles in both induction and repression of transcription. EMBO J. 5, 603–608. 10.1002/j.1460-2075.1986.tb04253.x.

74. Söding, J., Zwicker, D., Sohrabi-Jahromi, S., Boehning, M., and Kirschbaum, J. (2020). Mechanisms for Active Regulation of Biomolecular Condensates. Trends Cell Biol. 30, 4–14. 10.1016/j.tcb.2019.10.006.

75. Martin, E.W., and Holehouse, A.S. (2020). Intrinsically disordered protein regions and phase separation: sequence determinants of assembly or lack thereof. Emerg. Top. Life Sci. 4, 307–329. 10.1042/ETLS20190164.

76. Staller, M.V. (2022). Transcription factors perform a 2-step search of the nucleus. Genetics 222, iyac111. 10.1093/genetics/iyac111.

77. Gill, G., and Ptashne, M. (1988). Negative effect of the transcriptional activator GAL4. Nature 334, 721–724. 10.1038/334721a0.

78. Irgen-Gioro, S., Yoshida, S.R., Walling, V., and Chong, S. (2022). Fixation can change the appearance of phase separation in living cells. eLife 11, e79903. 10.7554/eLife.79903.

79. MacKinnon, J.G. (2007). Bootstrap Hypothesis Testing. Work. Pap.

