## Supplemental Information for "Transcription factor clusters enable target search but do not contribute to target gene activation"

### SUPPLEMENTAL FIGURES AND LEGENDS

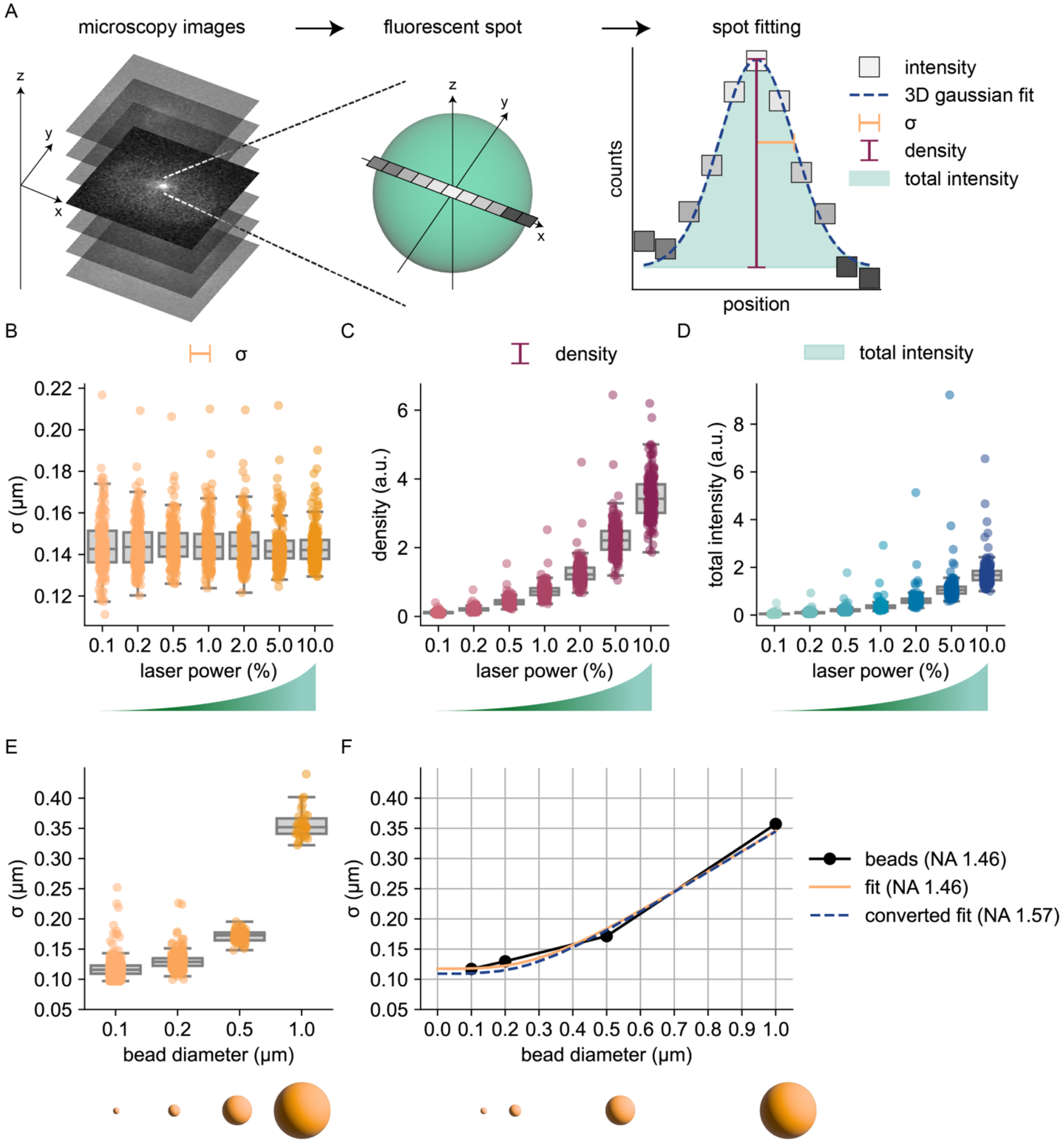


**Figure S1. Spot fitting algorithm to extract the cluster $\sigma$, density and total intensity.**

**A.** Schematic representation of data analysis, starting with a microscopy image of cells with fluorescent spots (clusters, DNA/RNA label), followed by a schematic of a detected spot with intensities measured at each pixel (grey squares). These intensities are fit to a 3D gaussian with background subtraction using a tilted plane, allowing extraction of $\sigma$ (the standard deviation, a measure for cluster size), cluster density (peak height, a measure for concentration within the cluster) and the total integrated intensity (as a measure for the total number of molecules in the cluster).

**B-D.** Distribution of $\sigma$, density and total intensity of 202 individual fluorescent beads (0.21 µm TetraSpec microspheres) measured at different laser powers representing the width, peak height, and integrated intensity of the 3D gaussian fit, respectively (see methods for details). As expected, the spot density, but not the $\sigma$ changes at different laser powers. Circles show data for individual beads and box plots show the distribution of the data, with box edges indicating first and third quartiles, center line indicating the median and whiskers indicating the 1.5x interquartile range.

**E.** Top: Distribution of $\sigma$ of individual fluorescent beads (Tetraspec microspheres) with increasing diameters (see methods for details). For every bead diameter multiple *z*-stacks were taken, resulting in a total of 676, 310, 69 and 33 detected beads of respectively 0.1, 0.2, 0.5 and 1.0 µm in diameter. As expected, the $\sigma$ changes for different bead sizes. Circles show data for individual beads and box plots show the distribution of the data, with box edges indicating first and third quartiles, center line indicating the median and whiskers indicating the 1.5x interquartile range. Bottom: schematic representation of increasing bead diameter (to scale).

**F.** Calibration curve to relate measured $\sigma$ to spot diameter. Black: mean measured values for $\sigma$ of the different diameter beads from E. Orange: fit to the black data taken with a NA 1.46 objective (see methods for details). Blue dashed line: conversion of fit curve to predicted relationship for NA 1.57 objective (as used for Gal4-EGFP clusters, see methods for details). Bottom: schematic representation of increasing bead diameter (to scale).

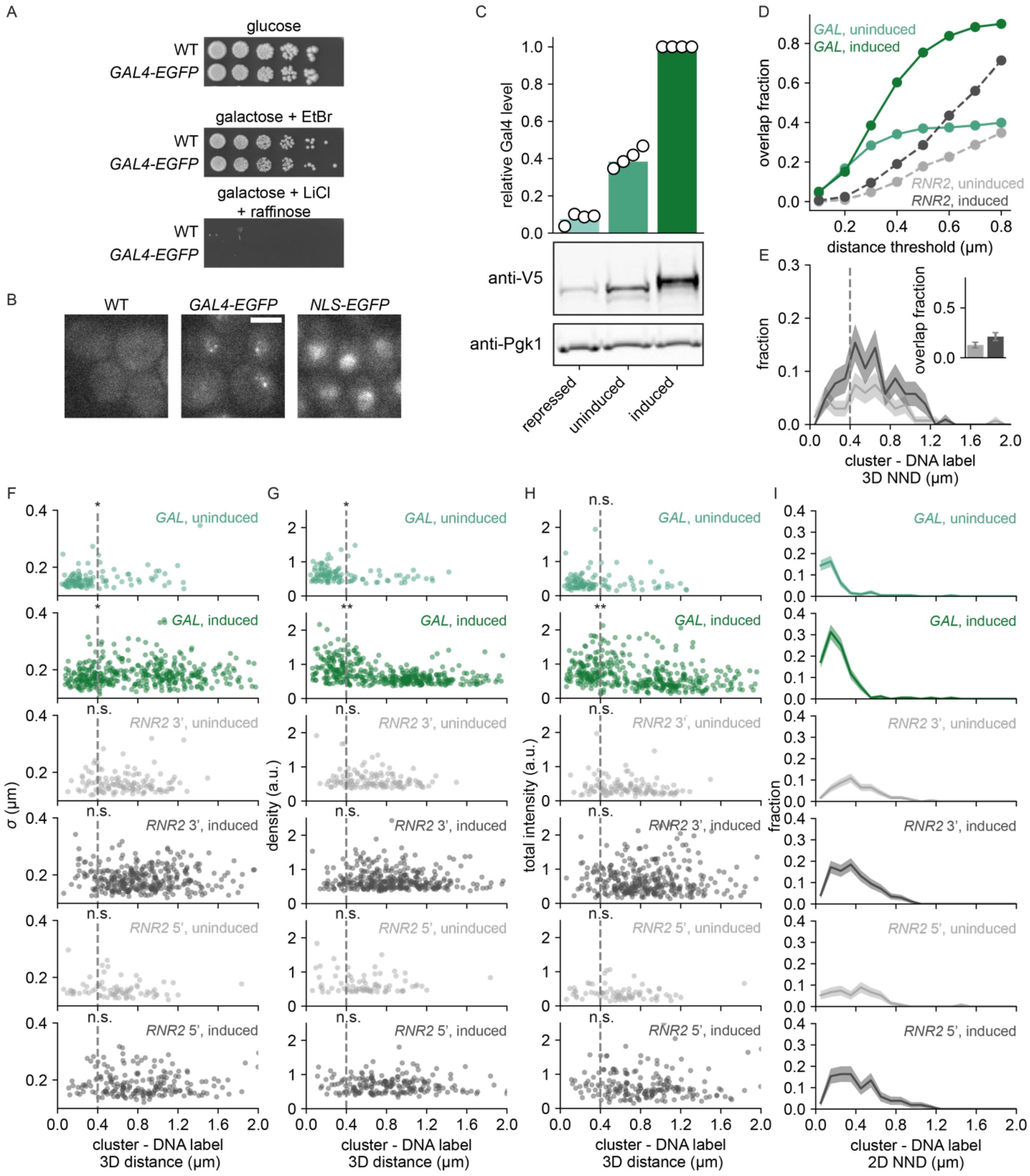


**Figure S2. Gal4 forms clusters that colocalize with the *GAL* genes. Related to Figure 1.**

**A.** Growth assay of indicated strains to assess their galactose metabolism capability. Shown are 5-fold serial dilutions on YEP + 2% glucose (dilution control), YEP + 2% galactose + 20 μg/mL ethidium bromide (growth = functional galactose metabolism) and YEP + 2% raffinose + 2% galactose + 40 mM lithium chloride + 0.003% methionine (no growth = functional galactose metabolism).

**B.** Representative images of WT yeast cells (left), cells expressing Gal4-EGFP (middle) and cells expressing NLS-EGFP in induced (galactose and raffinose) conditions. Only for Gal4-EGFP clustering is observed indicating the specificity of Gal4 for this clustering. Images are a single *z*-slice of a representative group of cells. Scalebar: 3 μm.
**C.** Western blot quantification of Gal4-EGFP-V5 protein levels using an anti-V5 antibody measured in repressed (glucose), uninduced (raffinose) or induced (galactose) conditions. Expression levels are normalized to Pgk1 and to the expression level in 2.00% galactose. Open circles represent the results of individual replicate experiments, green bars indicate their mean. Western blot images are a representative example of 4 independent experiments.
**D.** Fraction of cells with an overlapping cluster with the *GAL* DNA label in uninduced (raffinose, light green) and induced (raffinose and galactose, dark green) conditions or with the *RNR2* DNA label in uninduced (light grey) and induced (dark grey) conditions for varying distance thresholds used to discriminate overlapping and non-overlapping clusters. At all distance thresholds, the *GAL* DNA labels show more overlapping clusters than the negative control gene, in induced and uninduced conditions.
**E.** Distribution of 3D nearest neighbor distances (NNDs) between the *RNR2* 5' DNA label and the closest cluster in uninduced (light grey, 172 cells) and induced (dark grey, 118 cells) conditions. Shaded regions represent SEM based on 1000 bootstrap repeats. Vertical dashed line indicates 400 nm threshold used to discriminate between overlapping and non-overlapping clusters. Inset shows fraction of DNA-label containing cells with an overlapping cluster. Error bars represent SEM based on 1000 bootstrap repeats.
**F-H.** Scatterplot of **F.** cluster $\sigma$, **G.** density and **H.** total intensity versus 3D distance between the Gal4-EGFP cluster and the *GAL* DNA label in uninduced (light green, 306 cells) and induced (dark green, 200 cells) conditions, the *RNR2* 3' DNA label in uninduced (light grey, 276 cells) and induced (dark grey, 198 cells) conditions, and the *RNR2* 5' DNA label in uninduced (light grey, 172 cells) and induced (dark grey, 118 cells) conditions. Vertical dashed line indicates 400 nm threshold used to discriminate between overlapping and non-overlapping clusters. Significance between cells closer and further than 400 nm from the DNA label was determined by Mann-Whitney *U* test; n.s.: not significant; *: $p < 0.05$; **: $p < 0.01$.
**I.** Distribution of 2D nearest neighbor distances (NND) between DNA label and the closest cluster for the same dataset and conditions as **F-H.** Shaded regions represent SEM based on 1000 bootstrap repeats.

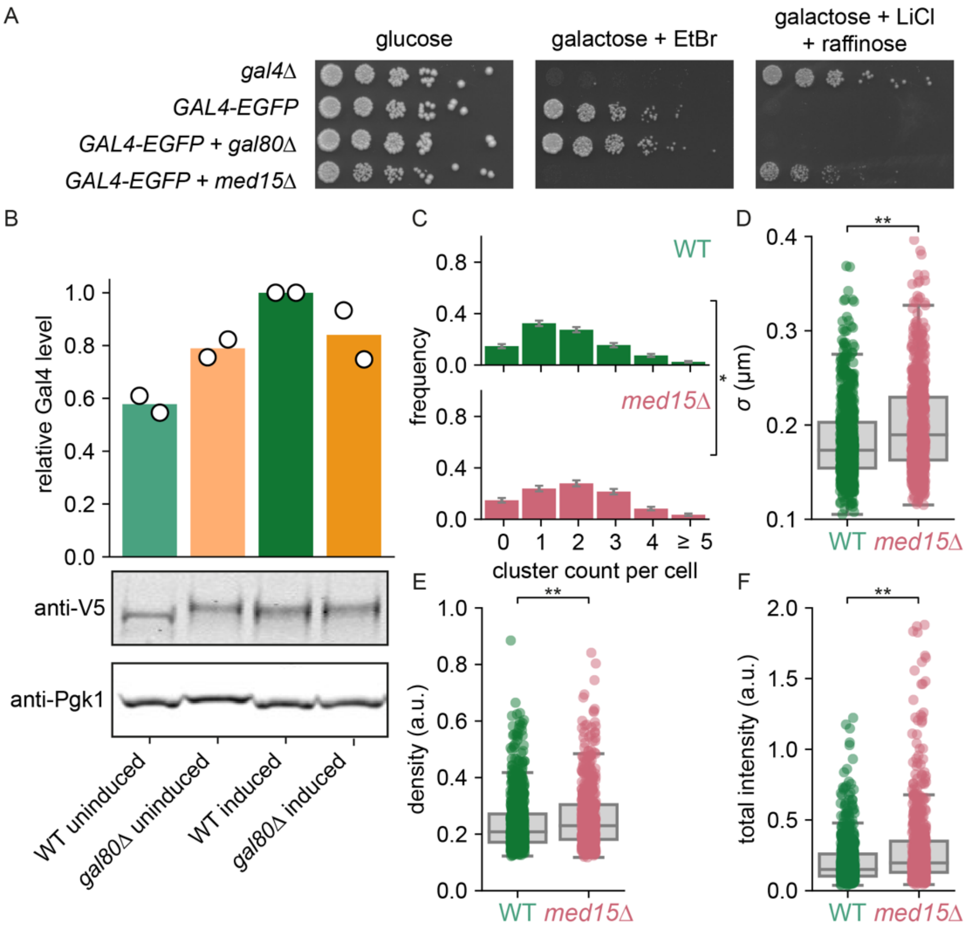


**Figure S3. Gal4 clusters are limited by Gal80 but do not depend on interactions with Med15. Related to Figure 2.**

**A.** Growth assay of indicated strains to assess their galactose metabolism capability. Shown are 5-fold serial dilutions on YEP + 2% glucose (dilution control), YEP + 2% galactose + 20 μg/mL ethidium bromide (growth = functional galactose metabolism) and YEP + 2% raffinose + 2% galactose + 40 mM lithium chloride + 0.003% methionine (no growth = functional galactose metabolism).

**B.** Western blot quantification of Gal4-EGFP-V5 protein levels using an anti-V5 antibody measured in WT and *gal80Δ* in uninduced (raffinose) and induced (galactose) conditions. Expression levels are normalized to Pgk1 and to the expression level of WT in induced conditions. Open circles represent the results of individual replicate experiments, green and orange bars indicate their mean. Western Blot images are a representative example of 2 independent experiments.

**C-F.** Quantification of Gal4-EGFP clusters in WT (green, 484 cells) and *med15Δ* (pink, 372 cells) in induced (raffinose and galactose) conditions. **C.** Distribution of number of clusters observed per cell. Error bars indicate SEM based on 1000 bootstrap repeats; *: $p<0.05$. **D-F.** Distribution of **D.** cluster $\sigma$, **E.** density and **F.** total intensity. Circles show data for individual clusters and box plots show the distribution of the data, with box edges indicating first and third quartiles, center line indicating the median and whiskers indicating the 1.5x interquartile range. Significance determined by Mann-Whitney *U* test; *: p<0.05; **: p < 0.01.

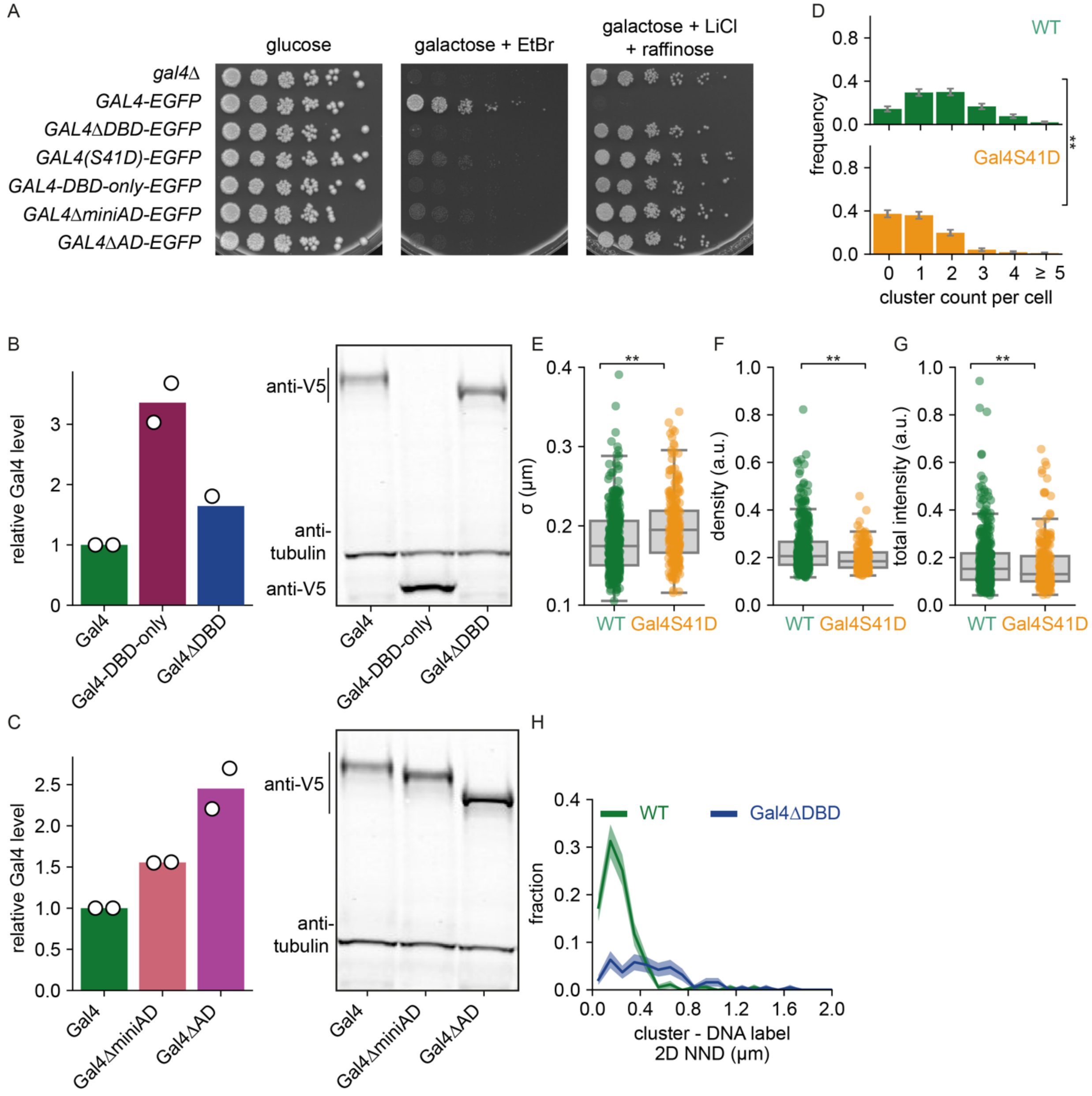


**Figure S4. Characterization of different Gal4 mutants. Related to Figure 3 and 4.**

**A.** Growth assay of indicated strains to assess their galactose metabolism capability. Shown are 5-fold serial dilutions on YEP + 2% glucose (dilution control), YEP + 2% galactose + 20 µg/mL ethidium bromide (growth = functional galactose metabolism) and YEP + 2% raffinose + 2% galactose + 40 mM lithium chloride + 0.003% methionine (no growth = functional galactose metabolism).

**B.** Left: western blot quantification of Gal4-EGFP-V5, Gal4-DBD-only-GFP-V5 and Gal4ΔDBD-EGFP-V5 protein levels using an anti-V5 antibody measured in induced (galactose and raffinose) conditions. Expression levels are normalized to tubulin and to the expression level of WT Gal4-EGFP-V5. Open circles represent the results of individual replicate experiments, colored bars indicate their mean. Right: corresponding western blot image, representative example of 2 independent experiments.

**C.** Left: western blot quantification of Gal4-EGFP-V5, Gal4ΔminiAD-GFP-V5 and Gal4ΔAD-GFP-V5 protein levels using an anti-V5 antibody measured in induced conditions. Expression levels are normalized to tubulin and to the expression level of WT Gal4-EGFP-V5. Open circles represent the results of individual replicate experiments, colored bars indicate their mean. Right: corresponding western blot image, representative example of 2 independent experiments.

**D-G.** Quantification of WT Gal4-EGFP (green, 210 cells) and Gal4(S41D)-EGFP (orange, 217 cells) clusters in induced conditions **D.** Distribution of number of clusters observed per cell. Error bars indicate SEM based on 1000 bootstrap repeats. **E-G.** Distribution of **E.** cluster $\sigma$, **F.** density and **G.** total intensity. Circles show data for individual clusters and box plots show the distribution of the data, with box edges indicating first and third quartiles, center line indicating the median and whiskers indicating the 1.5x interquartile range. Significance determined by Mann-Whitney *U* test; *: $p < 0.05$; **: $p < 0.01$.

**H.** Distribution of 2D nearest neighbor distances (2D NND) between the *GAL* DNA label and the closest WT Gal4-EGFP (green) or Gal4ΔDBD-EGFP (blue) cluster. Shaded regions represent SEM based on 1000 bootstrap repeats.

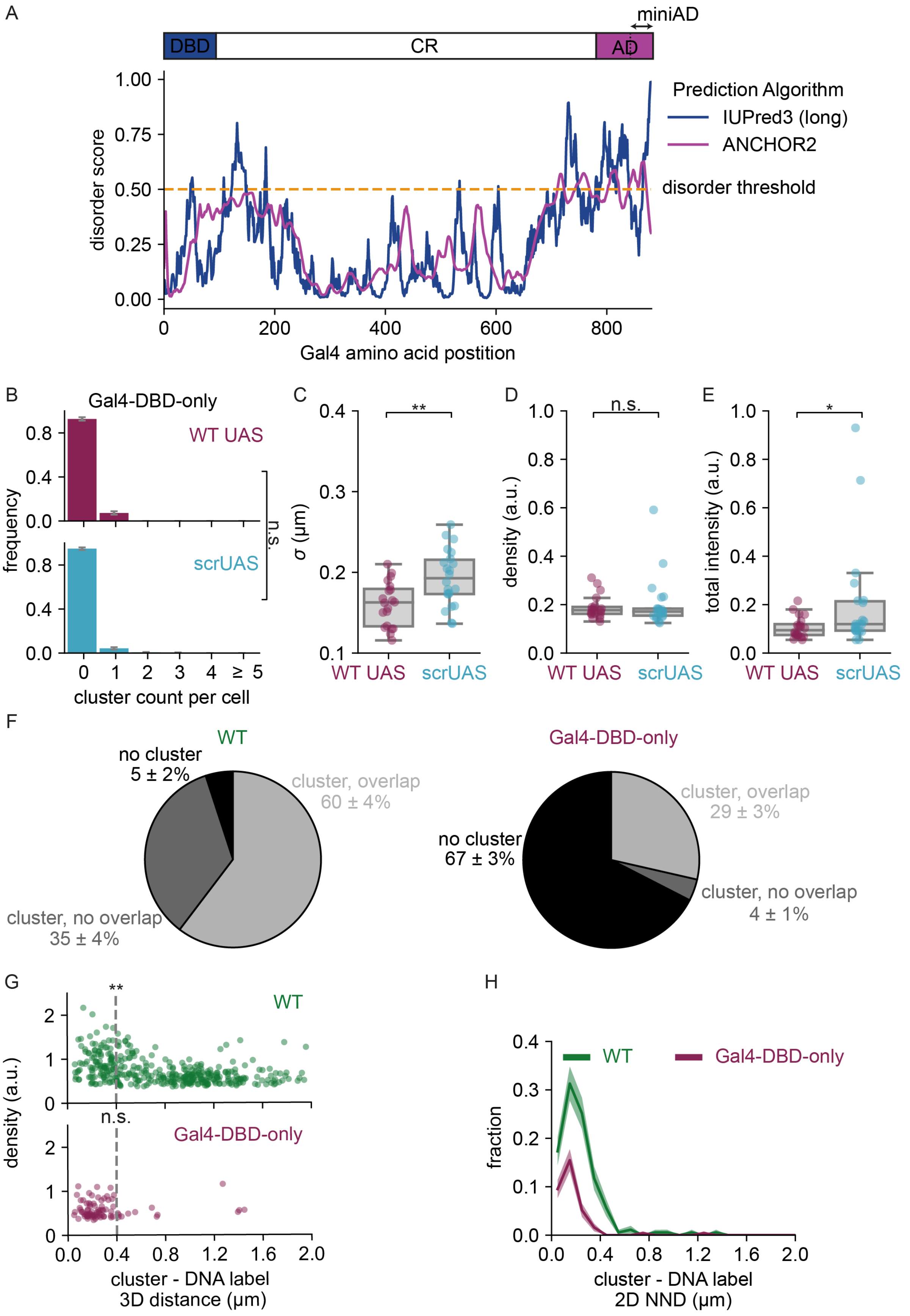
A
miniAD
DBD
CR
AD
Prediction Algorithm
IUPred3 (long)
ANCHOR2
disorder threshold
disorder score
Gal4 amino acid postition
B
Gal4-DBD-only
WT UAS
scrUAS
n.s.
frequency
cluster count per cell
C
σ (μm)
D
density (a.u.)
E
total intensity (a.u.)
WT UAS
scrUAS
F
WT
no cluster
5 ± 2%
cluster, overlap
60 ± 4%
cluster, no overlap
35 ± 4%
Gal4-DBD-only
cluster, overlap
29 ± 3%
no cluster
67 ± 3%
cluster, no overlap
4 ± 1%
G
WT
Gal4-DBD-only
density (a.u.)
cluster - DNA label
3D distance (μm)
H
WT
Gal4-DBD-only
fraction
cluster - DNA label
2D NND (μm)

**Figure S5. IDRs are not essential for clustering but contribute to target search. Related to Figure 4.**

**A.** Top: schematic representation of Gal4 protein domains as used in this study. Blue: DBD, white: region containing CR, purple: AD. Dashed line indicates miniAD domain. Bottom: Predicted disorder of the Gal4 protein based on IUPred3 long (blue) and ANCHOR2 (purple) prediction algorithms[1,2]. Orange dashed line indicates disorder threshold of 0.5 to discriminate ordered from disordered regions.

**B-E.** Quantification of Gal4-DBD-only-EGFP clusters in cells with WT UAS (purple, 274 cells) and scrUAS (cyan, 348 cells) in induced (galactose and raffinose) conditions. **B.** Distribution of number of clusters observed per cell. Error bars indicate SEM based on 1000 bootstrap repeats.

**C-E.** Distribution of **C.** cluster $\sigma$, **D.** density and **E.** total intensity. Circles show data for individual clusters and box plots show the distribution of the data, with box edges indicating first and third quartiles, center line indicating the median and whiskers indicating the 1.5x interquartile range. Significance determined by Mann-Whitney *U* test; n.s.: not significant; *: $p < 0.05$; **: $p < 0.01$.

**F.** Pie-charts showing the percentages of cells with a *GAL* DNA label that show a cluster overlapping with the *GAL* locus (light grey), a cluster that does not overlap with the *GAL* locus (dark grey), or do not show any clusters (black) for WT Gal4-EGFP (left, 179 cells) and Gal4-DBD-only-EGFP (right, 252 cells) in induced conditions. Error values are SEM based on 1000 bootstrap repeats.

**G.** Scatterplot of cluster density versus 3D distance between the cluster and the *GAL* DNA label for WT Gal4-EGFP (green, 200 cells) and Gal4-DBD-only-EGFP (purple 276, cells) in induced conditions. Vertical dashed line indicates 400 nm threshold used to discriminate between overlapping and non-overlapping clusters. Significance between clusters closer and further than 400 nm from the DNA label was determined by Mann-Whitney *U* test; n.s.: not significant; *: $p < 0.05$; **: $p < 0.01$.

**H.** Distribution of 2D nearest neighbor distances (2D NND) between the *GAL* DNA label and the closest cluster for WT Gal4 (green, 200 cells) and Gal4-DBD-only (purple, 276 cells), same dataset as in **G**. Shaded regions represent SEM based on 1000 bootstrap repeats.

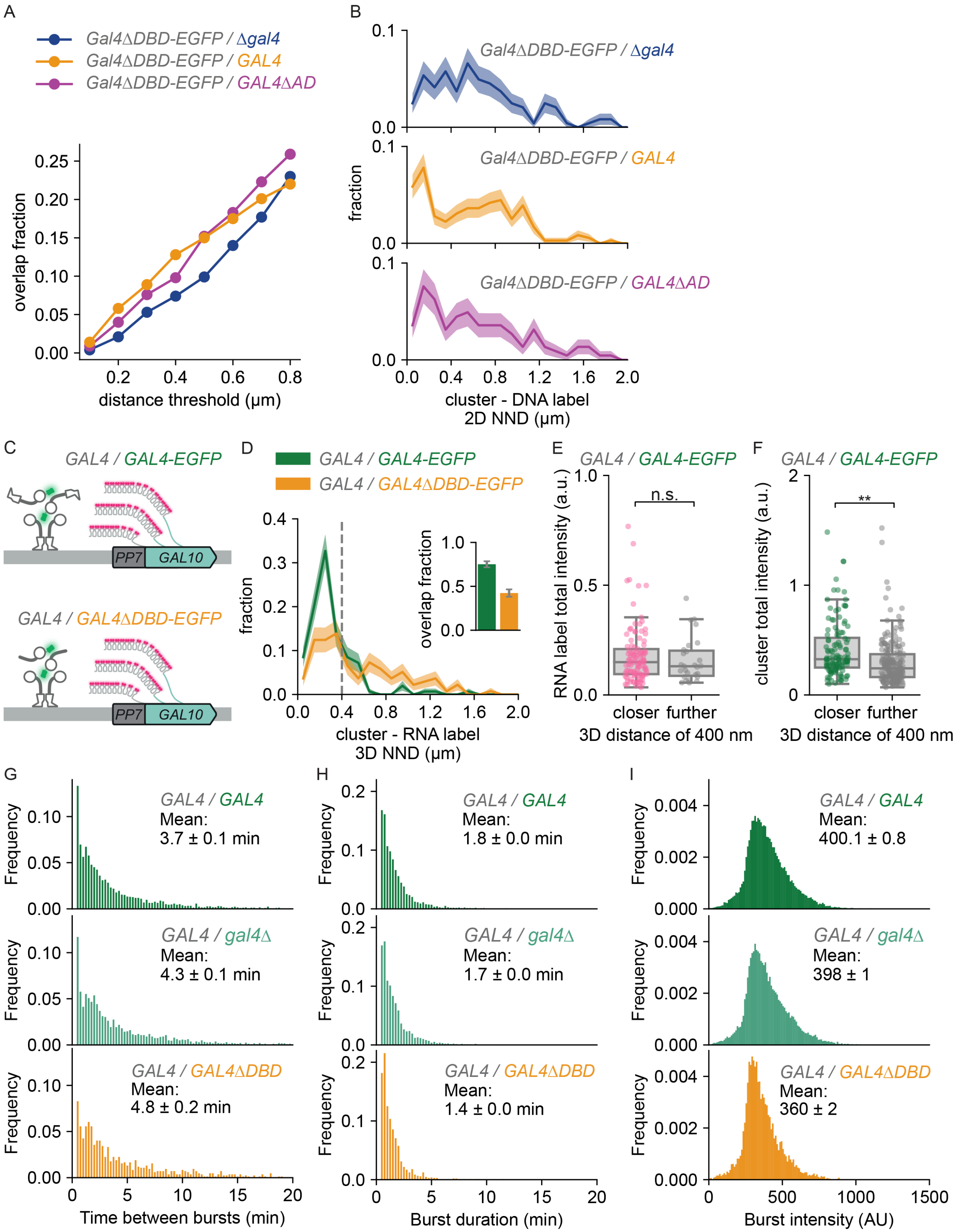
A
Gal4ΔDBD-EGFP / Δgal4
Gal4ΔDBD-EGFP / GAL4
Gal4ΔDBD-EGFP / GAL4ΔAD
overlap fraction
distance threshold (µm)
B
Gal4ΔDBD-EGFP / Δgal4
Gal4ΔDBD-EGFP / GAL4
Gal4ΔDBD-EGFP / GAL4ΔAD
fraction
cluster - DNA label
2D NND (µm)
C
GAL4 / GAL4-EGFP
GAL4 / GAL4ΔDBD-EGFP
PP7
GAL10
D
GAL4 / GAL4-EGFP
GAL4 / GAL4ΔDBD-EGFP
fraction
overlap fraction
cluster - RNA label
3D NND (µm)
E
GAL4 / GAL4-EGFP
RNA label total intensity (a.u.)
n.s.
closer further
3D distance of 400 nm
F
GAL4 / GAL4-EGFP
cluster total intensity (a.u.)
**
closer further
3D distance of 400 nm
G
GAL4 / GAL4
Mean:
3.7 ± 0.1 min
GAL4 / gal4Δ
Mean:
4.3 ± 0.1 min
GAL4 / GAL4ΔDBD
Mean:
4.8 ± 0.2 min
Frequency
Time between bursts (min)
H
GAL4 / GAL4
Mean:
1.8 ± 0.0 min
GAL4 / gal4Δ
Mean:
1.7 ± 0.0 min
GAL4 / GAL4ΔDBD
Mean:
1.4 ± 0.0 min
Frequency
Burst duration (min)
I
GAL4 / GAL4
Mean:
400.1 ± 0.8
GAL4 / gal4Δ
Mean:
398 ± 1
GAL4 / GAL4ΔDBD
Mean:
360 ± 2
Frequency
Burst intensity (AU)

**Figure S6. Gal4 self-interactions are sufficient to recruit Gal4 to target genes but insufficient to activate transcription. Related to Figure 5 and 6.**

**A.** Fraction of cells with a Gal4ΔDBD-EGFP cluster overlapping with the *GAL* DNA label for *GAL4ΔDBD-EGFP/gal4Δ* (blue, 300 cells), *GAL4ΔDBD-EGFP/GAL4* (orange, 423 cells) and *GAL4ΔDBD-EGFP/Gal4ΔAD* (purple, 256 cells) in induced conditions (raffinose and galactose) for varying distance thresholds used to discriminate between overlapping and non-overlapping clusters. Addition of WT Gal4 or Gal4ΔAD results in increased overlap of Gal4-ΔDBD clusters with the *GAL* DNA label regardless of the chosen threshold.
**B.** Distribution of 2D nearest neighbor distances (2D NND) between the *GAL* DNA label and the closest cluster for *GAL4ΔDBD-EGFP/gal4Δ* (blue, 300 cells), *GAL4ΔDBD-EGFP/GAL4* (orange, 423 cells) and *GAL4ΔDBD-EGFP/Gal4ΔAD* (purple, 256 cells) in induced conditions. Shaded regions represent SEM based on 1000 bootstrap repeats.
**C.** Schematic representation of the *PP7-GAL10* gene and diploid yeast strains expressing WT Gal4 from one allele and on the other allele expressing either WT Gal4-EGFP (*GAL4/GAL4-EGFP,* green) or Gal4ΔDBD-EGFP (*GAL4/GAL4ΔDBD-EGFP*, orange*)*. Nascent *GAL10* transcripts are fluorescently labeled by binding of the PP7-coat protein-ymScarletI to the PP7-stem loops (magenta), forming bright RNA label spots indicating the location of the *GAL10* transcription site and thereby the *GAL* locus*.*
**D.** Distribution of 3D nearest neighbor distances (NND) between the *PP7-GAL10* RNA label and the closest Gal4-EGFP (green, 161 cells) or Gal4ΔDBD-EGFP cluster (blue, 141 cells) in the presence of WT Gal4 in induced (raffinose and galactose) conditions. Shaded regions represent SEM based on 1000 bootstrap repeats. Vertical dashed line indicates 400 nm threshold used to discriminate between overlapping and non-overlapping clusters. Inset shows fraction of RNA label-containing cells with an overlapping cluster. Error bars represent SEM based on 1000 bootstrap repeats.
**E.** Distributions of the total intensity of the *GAL10* RNA labels in *GAL4/GAL4-EGFP* yeast cells (same dataset as **D.**) with a cluster closer than 400 nm of the label (pink, 105 cells) or further (grey, 27 cells). Circles show data for individual RNA labels and box plots show the distribution of the data, with box edges indicating first and third quartiles, center line indicating the median and whiskers indicating the 1.5x interquartile range. Significance determined by Mann-Whitney *U* test; n.s.: not significant.
**F.** Distributions of total intensity of the Gal4-EGFP clusters in *GAL4/GAL4-EGFP* yeast cells (161 cells, same dataset as **D.**) closer than 400 nm from the *GAL10* RNA label (green) or further (grey). Circles show data for individual clusters and box plots show the distribution of the data, with box edges indicating first and third quartiles, center line indicating the median and whiskers indicating the 1.5x interquartile range. Significance determined by Mann-Whitney *U* test; **: $p < 0.01$.
**G-I.** Distributions of **G.** the time between bursts, **H.** burst duration and **I.** burst intensity for *PP7-GAL10* transcription in *GAL4*/*GAL4* (green, 673 cells), *GAL4/gal4Δ* (light green, 669 cells) and *GAL4/GAL4ΔDBD* (orange, 265 cells) yeast strains. Values indicated in the graph are the mean with standard deviation based on 1000 bootstrap repeats.

MATERIALS & METHODS

#### Yeast strains and plasmids

All strains were derived from BY4741 and BY4742 parent strains. The BY4742 *GAL4-EGFP* strain (YTL390) was created by transformation using a PCR product with EGFP and *loxP-kanMX-loxP* followed by kanMX removal by CRE recombinase. The BY4742 *gal4Δ* strain (YTL559) was created by transformation using a PCR produced containing a kanMX cassette.

Truncations and mutations of *GAL4* were created using a CRISPR-Cas9-based approach[3]: *BPSV40-GAL4(Δ1-94)* (Gal4ΔDBD; YTL1284 and YTL1662), *GAL4(Δ840-881)* (Gal4ΔminiAD; YTL1221), *GAL4(Δ768-881)* (Gal4ΔAD; YTL1286), *GAL4(Δ95-881)* (Gal4-DBD-only; YTL1226, YTL1639 and YTL1686), *GAL4::BPSV40* (YTL1702) and GAL4(S41D) (YTL945). Strains were transformed using a plasmid expressing Cas9 and a guide RNA and either double-stranded PCR repair template or single-stranded oligo, followed by removal of the Cas9 plasmid by 5-FOA selection.

To scramble the Gal4 UAS sites (scrUAS) at *GAL2*, *GAL7* and *GAL10* (YTL1154), three successive rounds of transformations were used to edit one locus at a time using the CRISPR-based approach described above with single-stranded oligos as repair templates.

To introduce the DNA label at the *GAL* locus (*3'-GAL1*; in YTL1652, YTL1662 and YTL1686), *3'-RNR2* (in YTL1699) or *5'-RNR2* (YTL1698), three successive rounds of transformations were used. First, a natMX cassette was integrated at either *3'-GAL1*, *3'-RNR2* or *5'-RNR2* by transformation with a PCR product encoding for natMX and homology arms for the *tetO* array. Next, the natMX cassette was replaced with *tetOx128* array using the CRISPR-based approach described above using a PCR product as repair template. Finally, *tetR1-tdTomato* was integrated at the *ADE1* locus by transformation using a plasmid digestion as a repair template. The BY4743 strains with the DNA label at the *GAL* locus (YTL1678 and YTL1693) were created by mating the BY4742 strain with *GAL* DNA label and *GAL4-EGFP* or *BPSV40-GAL4(Δ1-94)-EGFP* with either a BY4741 Δ*gal4* strain (YTL1679), a WT BY4741 strain (YTL1678) or a BY4741 *Gal4(Δ768-881)* strain (YTL1693).

The BY4743 diploid strains with PP7 loops (YTL1218, YTL1317 for cluster-RNA-label overlap, YTL1098, YTL1326 for growth assay and YTL590, YTL1431 and YTL1432 for live-cell imaging) were created by mating of a BY4741 and a BY4742 haploid yeast strain. The BY4742 strain was either *GAL4-EGFP*, *BPSV40-GAL4(Δ1-94)-EGFP* or *gal4*Δ. The BY4741 strain contained 14xPP7-GAL10, inserted by transformation with a PCR product containing the PP7 loop cassette and *loxP-kanMX-loxP* followed by *kanMX* removal by CRE recombinase. The PP7 coat protein was inserted in the BY4741 strain by transformation with a digested plasmid as a repair template (pTL174:PacI for PCP-GFPEnvy in YTL590, YTL1431 and YTL1432 or pTL306:PacI for PCP-ymScarletI in YTL1317, YTL1326 and YTL1218).

The BY4742 with *GAL4-EGFP* + *gal80Δ* strain (YTL762) was created by transformation of the BY4742 with *GAL4-EGFP* strain with a PCR product containing a *loxP-kanMX-loxP* cassette to replace *GAL80* and subsequent *kanMX* removal by CRE recombinase. For the BY4742 with *GAL4-EGFP* and *med15Δ*, the *med15Δ* was created using the CRISPR-based approach described above using a single-stranded oligo as a repair template.

For Western Blot experiments, V5-tags (3x V5) were introduced using the CRISPR-based approach described above using a PCR product as a repair template, using YTL390, YTL762, YTL1284, YTL1221, YTL1286 and YTL1226 as parent strains for YTL1653, YTL1685, YTL1655, YTL1661 and YTL1654, respectively. For all strains at least two replicates were constructed independently, which were verified by PCR and, if applicable, sequencing.

All strains, plasmids and oligos used in this study are listed in Table S1, S2 and S3 respectively. Yeast strains and plasmids are available on request.

##### Live-cell imaging of Gal4 clustering

Yeast cultures were grown to early mid-log in synthetic complete medium and imaged on a coverslip with a 2% agarose pad, as described previously[4]. Both the synthetic complete medium and agarose pad contained the indicated carbon sources. For cells with a DNA-label, containing the *ade1::tetR1-tdTomato-kanMX* integration, 40 mg/L adenine was added to both synthetic complete medium and agarose pad to rescue *ade1* deficiency.

Imaging was performed on an AxioObserver.7 / ELYRA.P1 microscope (Zeiss) equipped with an incubator for microscopy (Pecon) set at 30ºC, Scanning Stage Piezo 130x100 (Zeiss) and 405, 488, 561 and 640 nm lasers (Coherent) with maximum powers at 50, 100, 100 and 150 mW respectively. We used an alpha Plan-Apochromat 100× NA 1.57 oil objective (Zeiss), and a filterset consisting of a ZT405/488/561rpcv2-UF1 dichroic filter (Chroma) and a ZET405/488/561/640mv2 emission filter (Chroma). The emission was split in two channels (TV1 and TV2) using a duolink splitter (Zeiss) holding a filterset with a BS561 dichroic beamsplitter (Zeiss) and FF03-525/50-25 and BLP02-561R-25 emission filters (Semrock) used for imaging DNA- or RNA-labels and Gal4-EGPF clusters respectively on two EM-CCD iXon DU 897 camera's (Andor). All imaging was performed using the following settings in Zen Blue software: TIRF acquisition mode, 512x512 pixels field of view, 1.6x optovar, HILO illumination mode, 50 ms exposure time, EMCCD gain set to 100× and *z*-stacks (using the piezo) were set to 21 planes at 250 nm intervals.

The Gal4-EGFP clusters were imaged with excitation at 488 nm at 25% power resulting in resulting in a ±2 kW/cm$^2$ excitation intensity. When imaging either the DNA-label or RNA-label, an extra *z*-stack (TV1) was taken prior to the *z*-stack capturing the clusters (TV2), using excitation at 561 nm at 0.2% power for the DNA label (±16 W/cm$^2$ excitation intensity) and 0.1% power for the RNA-label (±8 W/cm$^2$ excitation intensity).

Replicates of conditions to be compared were always imaged on the same day and comparisons were only made between these (paired-)replicates.

##### Image segmentation

For all experiments, the EGFP channel (TV2) was used for segmentation. Cells were detected and images were segmented accordingly using a custom Python script (https://github.com/Lenstralab/focianalysis). Briefly, a maximum intensity projection of the 3D z-stack was made which was then smoothed using a gaussian filter with $\sigma$ 2.5 voxels. The resulting image was then thresholded using Otsu's method to find area's containing cells. Any holes in this mask were filled and small features were removed. Finally, the cells were separated using watershedding with local maxima at least 40 pixels apart functioning as starting points. Then the area of the nucleus was determined for each cell individually by thresholding the cell. The threshold was determined by Otsu's method on the 75% of brightest non-zero pixels. The part of the cell higher than this threshold was taken to be the nucleus. After this automated segmentation, the cellular and nuclear masks were checked manually and corrected when multiple cells were masked together or when a mask contained debris or a dead cell.

##### Spot detection

Initial spot detection was done by applying a difference of gaussians filter (DoG) to the 3D $z$-stack. The DoG filter was applied with $\sigma$ 2.075 and 1.245 pixels in $x$ and $y$ directions and 1.325 and 0.795 planes in $z$ direction. The local maxima found were considered to be spot candidates.

Various absolute thresholds were used to select only local maxima above the background. These thresholds were kept the same for all conditions and replicates which were compared directly.

##### Spot fitting

A region of interest (ROI) of 11 pixels in $x$ and $y$ directions and 7 planes in $z$ direction was cut out of the image around each spot candidate. Then the parameters of this spot were determined in two steps: 1) iterative moment analysis and 2) fitting.

We assume each spot can be described by a gaussian profile on top of a tilted background:

$$G(x,y,z) = \frac{I}{(2\pi)^{3/2}\sigma_{xy}^2\sigma_z}\exp\left[-\frac{(x-x_0)^2+(y-y_0)^2}{2\sigma_{xy}^2}-\frac{(z-z_0)^2}{2\sigma_z^2}\right]+b+b_x x+b_y y+b_z z.$$

To ensure that we are calculating the parameters of the spot candidate instead of a nearby neighbor, the image in the ROI is multiplied by a gaussian approximating the microscope point spread function (psf, $\sigma_{xy}$ = 1.66 pixels and $\sigma_z$ = 1.06 pixels), centered in each iteration on the location $x_0, y_0, z_0$ of the spot determined by moment analysis in the step before. Besides the location, the background $b, b_x, b_y, b_z$ is determined in each iteration by fitting linear functions through the voxels at the edges of the ROI. This process is stopped when either the location does not significantly improve anymore between iterations, when the new location is more than 3 voxels from the spot candidate location or when 20 or more iterations were performed. Thereafter, the spot intensity $I$ and width $\sigma_{xy}, \sigma_z$ are determined by moment analysis, correcting for the limited domain of the ROI as moment analysis expects an infinite domain. The resulting parameters were then used as an initial guess for an Limited-memory Broyden-Fletcher-Goldfarb-Shanno (L-

BFGS-B) optimization on the sum of weighted squared log residuals, again to make sure that the spot candidate is fitted. The weight is defined as a gaussian with $\sigma_{xy}$ = 3.32 pixels and $\sigma_z$ = 2.12 pixels centered on the last position determined by the moment analysis. Subsequently, the goodness of fit is determined as the adjusted coefficient of determination for both the fit as a whole and for the peak and background parts individually. Finally, the peak intensity $I_p$, defined as the height of the maximum of the gaussian fit above the background, was calculated as $I_p = \frac{I}{(2\pi)^{3/2}\sigma_{xy}^2\sigma_z}$.

##### Spot filtering

Spots of which the resultant location was not within 3 voxels from the initial guess were considered to have a failed localization and removed from the results. Additionally, of peaks closer to each other than 0.1 times the point-spread-function size ($\sigma_{xy}$ = 1.66 pixels and $\sigma_z$ = 1.06 pixels), only the first was kept. Finally, only spots residing within the cellular masks and of which the goodness of fit (adjusted $R^2$) of the peak was above -1 were taken into account in the analysis of clusters and DNA-/RNA-labels.

##### Quantification of clusters

The frequency of number of clusters per cell was quantified by counting the number of filtered spots within every cell mask, combining the counts of every replicate per condition and thereafter making a normalized histogram of these counts in which the error bars represent the bootstrapped standard error of the mean (1000 repeats).

To compare cluster $\sigma$, density ($I_p$) and total intensity ($I$) between conditions, the filtered spots of all replicates within a condition were combined and were shown using boxplots in which the box indicates the quartiles of the dataset and the whiskers extend to 1.5 times the inter-quartile range. The boxplots were overlayed with the individual data points. For visualization purposes, the axis range was chosen such that it was easy to compare the boxplots between conditions. Note that in some cases, this led to a few single datapoints ending up outside of the displayed range of the plots. We note that all values, also those outside the displayed range, were included during statistical testing.

For both the number of clusters per cell, cluster $\sigma$, density and total intensity, the differences in population median between multiple conditions were first tested using the Kruskal-Wallis H-test for independent samples, followed by 2-sided pairwise Mann-Whitney *U* tests.

##### Quantification of colocalization between clusters and the DNA-/RNA-label

On every day of two-channel imaging, a *z*-stack of 0.21 µm TetraSpeck™ microspheres (ThermoFisher) was made to correct for aberrations between the channels of the DNA-/RNA-label (hereafter named 'reference label'; TV1) and clusters (TV2). In brief, a 2D affine transformation mapping TV2 to TV1 was determined using SimpleElastix and max *z* projections of both channels of the two-color bead sample, correcting for aberrations in *x* and *y*. For *z*, we assumed a simple (focus) offset, which was determined by processing the bead sample using the same spot detection and fitting pipeline as for the clusters, with the minor modification of using a 10x standard deviation threshold for determining the local maxima. We then found the nearest neighbor pairs between channels. These pairs were filtered from outliers using the interquartile range rule and the offset in the *z* direction was determined as the mean distance between the beads in each pair in *z*. Finally, the locations of the spots in the second channel (TV2) where corrected using the affine transformation and the offset in *z* prior to distance calculations.

After spot filtering, the reference label of every cell was determined as the spot with the largest density ($I_p$) within the nuclear mask, as both the DNA- and RNA-labels are expected to be in the nucleus. Within each reference label-containing cell, all 3D and 2D cluster-reference distances were calculated. The distributions of the beforementioned spot parameters ($\sigma$, density and total intensity) of spots with a distance closer or further than the overlap threshold were tested for differences using the 2-sided Mann-Whitney *U* test.

For the calculation of fraction of nearest neighbor distances (NNDs) all cells with a reference label were taken into account. For every cell, the NND was determined as the minimal 3D/2D distance. Cells with a reference label but lacking clusters were also included as fractions were normalized to all cells containing a reference label, but their NNDs were set to 'nan'. Finally, the NNDs were divided over 0.1 µm bins in a histogram normalized to all cells with a reference label. The standard error of the mean of every bin was calculated using bootstrapping (1000 repeats). Differences between conditions in populations of cells with a overlapping cluster-reference label were calculated using a 2-sided Fisher's exact test.

**Quantification of $\sigma$ and density are independent**

To test whether the spot fitting algorithm can independently fit spot $\sigma$ and density, *z*-stacks were taken of fluorescent beads (0.21 µm TetraSpeck microspheres, ThermoFisher)) at different laser powers. Apart from the varying laser powers, imaging settings were as before. Spots were localized with a standard deviation threshold of 10 for the detection of local maxima. Spots for which localization failed were filtered out as before. As expected, the $\sigma$ stays constant while the spot density increases changes at different laser powers (Figure S1B-D).

**Fit of $\sigma$ versus diameter**

To estimate the relationship between the measured $\sigma$ and the spot diameters, *z*-stacks were taken of 100, 200, 500 and 1000 nm fluorescent beads (TetraSpeck™ microspheres, Invitrogen T14792). Imaging settings were as before, with the minor modification that an alpha Plan-Apochromat 100× NA 1.46 oil objective was used as this corresponded to the coverslip on which the beads were mounted. Spots were localized with an adjusted DoG filter for NA 1.46 and a standard deviation threshold of 10 for the detection of local maxima. Spots for which localization failed were filtered out as before. The relationship between the $\sigma$ (Figure S1E), and the diameter $d$ was described empirically as:

$$d = \left(\frac{\sigma^{\beta} - \sigma_0^{\beta}}{\alpha}\right)^{\frac{1}{\beta}}$$

Fitting this equation to the bead diameters and the mean $\sigma$ for each bead diameter resulted in $\sigma_0 = 0.118 \pm 0.014$ µm, $\alpha = 0.05 \pm 0.05$ µm and $\beta = 3.46 \pm 2.15$. To convert the NA 1.46 results to NA 1.57, the acquired $\sigma_0$ was multiplied with the ratio 1.46/1.57, giving $\sigma_0 = 0.109 \pm 0.012$ µm (Figure S1F). Using these parameters, we converted our measurements of $\sigma$ for Gal4-EGFP clusters in induced (galactose) conditions (Figure 1D) to estimate the cluster diameter to be in the range of 100-763 nm. For this conversion, we used the 1.5x interquartile range as minimum and maximum values for $\sigma$.

**Live-cell imaging of transcription dynamics**

Live-cell imaging of transcription dynamics was performed as previously described in detail with minor modifications[4–6]. In brief, cells were imaged at mid-log ($OD600_{nm}$ 0.2-0.4) on a coverslip with an agarose pad consisting of 2% agarose and synthetic complete medium containing 2% galactose and 2% raffinose. Imaging was performed on a setup consisting of an AxioObserver inverted microscope (Zeiss), an alpha Plan-Apochromat 100× NA 1.46 oil objective, an sCMOS ORCA Flash 4v3 (Hamamatsu) with a 475-570 nm dichroic (Chroma), 570 nm longpass beamsplitter (Chroma) and 515/30 nm emission filter (Semrock), a UNO Top stage incubator (OKOlab) at 30 C, and LED excitation at 470/24 nm (SpectraX, Lumencor) at 20% power and an ND 2.0 filter, resulting in a 62 mW/cm$^2$ excitation intensity. Widefield images were recorded for one hour at 15 s interval, with *z*-stacks (9 slices, Dz 0.5 µm) and 150 ms exposure using Micro-Manager software[7]. For each condition, 9 replicate datasets were acquired with in total at least 265 cells.

**Analysis of transcription dynamics**

For analysis of the transcription dynamics imaging data, a similar approach was used as described previously[5]. All analysis was implemented as custom-written Python software (https://github.com/Lenstralab/livecell_analysis). First, images were corrected for *xy*-drift in the stage using an affine transformation on the maximum intensity projection. Next, cells were segmented using Otsu thresholding and watershedding. The intensity of the transcription sites (TS) was calculated by fitting a 2D Gaussian mask after local background subtractions as described previously[8]. Initially, a threshold of eight times the standard deviation of the background was used. For frames where no TS was detected, a second fit was made in the vicinity of the high intensity spots detected in that cell, using a threshold of six times the standard deviation of the background. For frames where no TS was detected after this second fit, the intensity was measured at the location of the previous frame where a TS was successfully found. The tracking within each cell was inspected visually, and the endpoint of each trace was manually set at the last frame where a TS is visible. Cells without a TS, dividing cells, cells that were segmented incorrectly and cells that contained tracking errors were excluded from analysis.

To determine the on and off periods, binarization was performed using a threshold set at six times the standard deviation of the background. The standard deviation of the background was determined for each cell fitting by a Lorentzian distribution to intensities measured at four points at a fixed distance from the TS in each frame in the same cell. This threshold was chosen to reliably distinguish on and off periods from background levels at the single-transcript level. Subsequently, the binarization was improved by removing bursts that last a single frame and merging bursts that are separated by a single frame. From these binarized traces, the burst durations, time between bursts, induction time are directly calculated. The burst intensity is measured as the average intensity of all frames in which the cell was on. The fraction of active cells was determined by manual scoring of the cell that do and the cells that do not show a TS during the one hour acquisition period.

In total at least 265 cells were included for each condition, and values for burst duration, time between bursts, induction time and burst intensity are determined by bootstrapping with 1000 repetitions. Reported error bars are standard deviations from the same bootstrap. Error bars in the number of active and inactive cells are given by the square root of the number of cells, as cells are expected to be independent of each other and thus follow Poisson statistics. To determine whether the obtained bursting parameters are significantly different between conditions, we have used bootstrap hypothesis testing using equation (4) from [9] to determine the achieved significance level.

**Protein detection by immunoblot and antibodies**

Yeast cultures were grown to $OD_{600nm}$ 0.5 in 25 mL synthetic complete media with indicated carbon sources, washed in MilliQ, pelleted and snap-frozen on dry ice. For protein extraction, cells were resuspended in 300 µl MilliQ, incubated with 300 µl 0.2M NaOH for 7 min at room temperature, centrifuged and resuspended in 500 µl 2× SDS-PAGE sample buffer (4% SDS, 20% glycerol, 0.1 M DTT, 0.125 M Tris-HCl pH 7.5 and EDTA-free protease inhibitors). Samples were incubated at 95°C for 5 min while shaking and centrifuged at 800g for 10 min at 4°C. A total of 20 µl lysate with loading buffer was run on a NuPAGE 3-8% gradient TAC gel and transferred to a 0.45-µm nitrocellulose membrane at 200 V, 1 A for 4h at 4°C. For blocking, the membrane was washed with TBS-T, incubated with PBS containing 5% milk for 1h and washed briefly with TBS-T, all at room temperature. The membrane was incubated with PBS containing 2% milk and primary antibody (1:5000) overnight at 4°C, washed three times with TBS-T for 10 min, incubated with 2% milk and secondary antibody (1:5000) for 1h at room temperature, washed three times with TBS-T for 10 min and once with PBS for 10 min, and imaged using an LI_COR Odyssey IR imager (Biosciences). Western blot analysis was performed using primary antibodies against V5 (R960-25, ThermoFisher), Pgk1 (Invitrogen 22C5D8, RRID: AB_2532235) and tubulin (Ab6161, Abcam) and secondary antibodies Odyssey goat-anti-mouse 800 nm and Odyssey goat-anti-rat 800 nm.

The fluorescence signal of western blot images was quantified using ImageJ[10,11]. In brief, ROIs of the same dimensions are drawn in each lane of the image. Next, a profile plot is created for each lane and a baseline is drawn manually to enclose the peak. The total area of the enclosed peak is calculated and used as a measure for the band intensity. This procedure is repeated for the signal of each primary antibody. The V5 band intensities are then normalized over corresponding Pgk1 or Tubulin bands and represented relative to the condition indicated in the figure legend.

**Growth assay**

The galactose metabolism capacity of yeast strains was assessed with a growth assay as described previously[5], with minor modifications. Serial five-fold dilutions of indicated strains (YTL559, YTL1284, YTL1286, YTL1326 and YTL1098 in Figure 6A; BY4742 and YTL390 in Figure S2A; YTL559, YTL390, YTL762 and YTL1304 in Figure S3A; YTL559, YTL390, YTL1284, YTL945, YTL1226, YTL1221 and YTL1286 in Figure S4A) were spotted on various plates and growth was assessed after 3 days at 30°C. Growth on YEP + 2% glucose was used as loading control. Growth on YEP + 2% galactose + 20 µg/mL ethidium bromide was interpreted as functional galactose metabolism, as galactose is the only carbon source available because ethidium bromide inhibits the use of amino acids as carbon source by binding to mitochondrial DNA. On the contrary, growth on YEP + 2% raffinose + 2% galactose + 40 mM lithium chloride (LiCl) + 0.003% methionine was interpreted as no functional galactose metabolism. Although galactose is present, its metabolism is lethal in the presence of LiCl due to the buildup of toxic metabolic intermediates. Therefore, only yeast without a functional galactose metabolism can survive on these plates, using the raffinose as carbon source. Methionine is added to prevent buildup of other toxic intermediates caused by LiCl inhibiting Hal2p/Met22p, the yeast BPNase[12].

**Data availability**

The microscopy data generated during the current study are available from the corresponding author on reasonable request. Software for analysis of clustering microscopy data is available at https://github.com/Lenstralab/focianalysis. Software code for analysis of transcription dynamics microscopy data is available at https://github.com/Lenstralab/livecell_analysis.

Supplementary Table 1: Yeast strains used in this study

| Strain | Genotype | Source |
|---|---|---|
| BY4741 | MATa *his3Δ1 leu2Δ0 met15Δ0 ura3Δ0* | Euroscarf |
| BY4742 | MATα *his3Δ1 leu2Δ0 lys2Δ0 ura3Δ0* | Euroscarf |
| BY4743 | MATa/α *his3Δ1/his3Δ1 leu2Δ0/leu2Δ0 LYS2/lys2Δ0 met15Δ0/MET15 ura3Δ0/ura3Δ0* | Euroscarf |
| YTL390 | BY4742 with *GAL4-EGFP* | This study |
| YTL1702 | BY4742 with *GAL4::BPSV40-EGFP* | This study |
| YTL762 | BY4742 with *GAL4-EGFP* + *gal80Δ* | This study |
| YTL1304 | BY4742 with *GAL4-EGFP* + *med15Δ* | This study |
| YTL1284 | BY4742 with *BPSV40-GAL4(Δ1-94)-EGFP* | This study |
| YTL945 | BY4742 with *GAL4(S41D)-EGFP* | This study |
| YTL1221 | BY4742 with *GAL4(Δ840-881)-EGFP* | This study |
| YTL1286 | BY4742 with *GAL4(Δ768-881)-EGFP* | This study |
| YTL1226 | BY4742 with *GAL4(Δ95-881)-EGFP* | This study |

| | | |
|---|---|---|
| YTL1653 | BY4742 with *GAL4-EGFP-3xV5* | This study |
| YTL1685 | BY4742 with *GAL4-EGFP-3xV5* + *gal80Δ* | This study |
| YTL1655 | BY4742 with *BPSV40-GAL4(Δ1-94)-EGFP-3xV5* | This study |
| YTL1661 | BY4742 with *GAL4(Δ840-881)-EGFP-3xV5* | This study |
| YTL1654 | BY4742 with *GAL4(Δ95-881)-EGFP-3xV5* | This study |
| YTL559 | BY4742 with *gal4Δ* | This study |
| YTL1154 | BY4742 with *scrUAS 5'GAL2* + *scrUAS 5'GAL7* + 3x *scrUAS 5'GAL10* + *GAL4-EGFP* | This study |
| YTL1639 | BY4742 with *scrUAS 5'GAL2* + *scrUAS 5'GAL7* + 3x *scrUAS 5'GAL10* + *GAL4(Δ95-881)-EGFP* | This study |
| YTL1652 | BY4742 with *tetOx128 3'-GAL1* + *ade1::tetR1-tdTomato-kanMX* + *GAL4-EGFP* | This study |
| YTL1698 | BY4742 with *tetOx128 5'-RNR2* + *ade1::tetR1-tdTomato-kanMX* + *GAL4-EGFP* | This study |
| YTL1699 | BY4742 with *tetOx128 3'-RNR2* + *ade1::tetR1-tdTomato-kanMX* + *GAL4-EGFP* | This study |
| YTL1662 | BY4742 with *tetOx128 3'-GAL1* + *ade1::tetR1-tdTomato-kanMX* + *BPSV40-GAL4(Δ1-94)-EGFP* | This study |
| YTL1686 | BY4742 with *tetOx128 3'-GAL1* + *ade1::tetR1-tdTomato-kanMX* + *GAL4(Δ95-881)-EGFP* | This study |
| YTL1679 | BY4743 with *GAL1/tetOx128 3'-GAL1* + *ADE1/ade1::tetR1-tdTomato-kanMX* + *BPSV40-GAL4(Δ1-94)-EGFP/gal4Δ* | This study |
| YTL1678 | BY4743 with *GAL1/tetOx128 3'-GAL1* + *ADE1/ade1::tetR1-tdTomato-kanMX* + *BPSV40-GAL4(Δ1-94)-EGFP/GAL4* | This study |
| YTL1693 | BY4743 with *GAL1/tetOx128 3'-GAL1* + *ADE1/ade1::tetR1-tdTomato-kanMX* + *BPSV40-GAL4(Δ1-94)-EGFP/GAL4(Δ768-881)* | This study |
| YTL1218 | BY4743 with *GAL10/14xPP7-GAL10* + *ura3Δ0/ura3Δ0::pRPL15A-PCP-ymScarletI* + *GAL4/GAL4-EGFP* | This study |
| YTL1317 | BY4743 with *GAL10/14xPP7-GAL10* + *ura3Δ0/ura3Δ0::pRPL15A-PCP-ymScarletI* + *GAL4/BPSV40-GAL4(Δ1-94)-EGFP* | This study |
| YTL1098 | BY4743 with *GAL10/14xPP7-GAL10* + *ura3Δ0/ura3Δ0::pRPL15A-PCP-ymScarletI* + *GAL4-EGFP/GAL4-EGFP* | This study |
| YTL1326 | BY4743 with *GAL10/14xPP7-GAL10* + *ura3Δ0/ura3Δ0::pRPL15A-PCP-ymScarletI* + *GAL4(Δ768-881)-EGFP/BPSV40-GAL4(Δ1-94)-EGFP* | This study |
| YTL590 | BY4743 with *GAL10/14xPP7-GAL10* + *ura3Δ0/ura3Δ0::pRPL15A-PCP-GFPEnvy* | This study |
| YTL1431 | BY4743 with *GAL10/14xPP7-GAL10* + *ura3Δ0/ura3Δ0::pRPL15A-PCP-GFPEnvy* + *GAL4/gal4Δ* | This study |
| YTL1432 | BY4743 with *GAL10/14xPP7-GAL10* + *ura3Δ0/ura3Δ0::pRPL15A-PCP-GFPEnvy* + *GAL4/BPSV40-GAL4(Δ1-94)* | This study |

Supplementary Table 2: Plasmids used in this study

| Plasmid name | Source |
|---|---|
| pTL014: pURA pGAL CRE recombinase (Euroscarf pSH47) | [13] |
| pTL031: 14x PP7 with loxP-kanMX-loxP (Addgene 189939) | [5] |
| pTL071: 1xEGFP tagging vector with loxP-kanMX-loxP | This study |
| pTL131: pML104 Cas9 guide RNA construct (Addgene 67638) | [3] |
| pTL174: pURA SIV pRPL15A PCP-NLS-EGFPEnvy (Addgene 189941) | This study |
| pTL306: pURA SIV pRPL15A PCP-NLS-ymScarlet-I | This study |
| pTL314: pML104 with guideRNA to GAL4 (ggcgcacttcggtttttctt) | This study |
| pTL317: pML104 with guideRNA to GAL4 (attcattttactcttttttt) | This study |
| pTL350: pML104 with guideRNA to GAL4 (gctactctcccaaaaccaaa) | This study |
| pTL354: pML104 with guideRNA to pGAL2 (ccgcacggacgaaagaccgc) | This study |
| pTL355: pML104 with guideRNA to pGAL7 (ttcggagcactgttgagcga) | This study |
| pTL363: pML104 with guideRNA to GAL4 (ggtgaaggccctactgagcc) | [6] |
| pTL370: pURA pADE3 BPSV40-EGFP | This study |
| pTL387: pML104 with guideRNA to GAL4 (atatacatcatccattgtag) | This study |
| pTL406: pML104 with gRNA to MED15 (taccctcggcacatttttct) | This study |
| pTL433: pML104 with gRNA to EGFP (ggatgaattgtacaaataac) | This study |
| pTL536: pDD207 tetR1-tdTomato | [14] |
| pTL539: pDD220 NAT-tet | [14] |
| pTL541: pDD226 tetOx128 | [14] |
| pTL551: pML104 with guideRNA to pTEF (gagattttgactgcaatttc) | This study |

Supplementary Table 3: Oligos used in this study (ordered from IDT)

| Oligo name | Sequence |
|---|---|
| GAL4-del-F | tgggacttgaacagctcctt |
| GAL4-del-R | tttgggtgtcttcatcacca |
| GAL4-GFP-F | gatgatgtatataactatctattcgatgatgaagataccccaccaaacccaaaaaa gaggccgctctagaactagtggatcc |

| | |
|---|---|
| GAL4-GFP-R | atggtgcacgatgcacagttgaagtgaacttgcggggtttttcagtatctacgattcatta gctgggtaccgcataggccac |
| GAL10-14PP7-5'-F | tattaaacttctttgcgtccatccaaaaaaaaagtaagaatttttgaaaattcaatataac aaagtgggagcgaggagatcc |
| GAL10-5'-R | agcaccacctgtaaccaaaacaattttagaagtactttcactttgtaactgagctgtcat gcataggccactagtggatctg |
| GAL4UAS_construct1 | aaaaccttctctttggaactttcagtaatacgcttaactgctcattgctatattgaagtaag ccgacccagagggttaggagccgccgagccgctcacggaagactctcctccgtgc gtcctcgtcttcaccggtcgcgttcctgaaacgcagatgtgcaccgtgcctcccccgtg cgaacaataaagattctacaatactagcttttatggttatgaagaggaaaaattggca gtaa |
| GAL4(S41>D) | gctccaaagaaaaaccgaagtgcgccaagtgtctgaagaacaactgggagtgtcg ctacgatcccaaaaccaaaaggtctccgctgactagggcacatctgacagaagtgg aatcaaggc |
| pGAL2_UAS2_repair | gatctatattcgaaaggggcggttgcctcaggaaggcactggctccttggggcctctg cggagatatctgcgccgttcaggggtccatgtgccttggacg |
| pGAL7_UAS2_repair | ttcttaaattgctttgcctctccttttggaaagctatacttgcagtccatggagggtcaagg ctcattagatatattttctgtcattttccttaacccaa |
| pGAL4_BPSV40-EGFP_F | gtgtctacgtaatgcacgccatcattttaagagaggacagagaagcaagcctcctga aagatgaaaaggacagcagatgg |
| pGAL4_BPSV40-EGFP_R | atggtgcacgatgcacagttgaagtgaacttgcggggtttttcagtatctacgattcatttt atttgtacaattcatccataccatgg |
| GAL4(del95-839)_insert | acatgattttgaaaatggattctttacaggatataaaagcattgttaacatggacggacc aaactgcgtataacgcgtttggaatcactacagggatgtt |
| GAL4(del840-881)-linker-EGFP_insert | cggctagtaaaattgatgatggtaataattcaaaaccactgtcacctggtgccgctcta gaactagtggatccgctgcaggaattcgatatcgtgtctaa |
| GAL4(del95-881)-linker-EGFP_insert | acatgattttgaaaatggattctttacaggatataaaagcattgttaacagccgctctag aactagtggatccgctgcaggaattcgatatcgtgtctaa |
| GAL4(del2-94)-EGFP | gcacgccatcattttaagagaggacagagaagcaagcctcctgaaagatgggatta tttgtacaagataatgtgaataaagatgccgtcacagatagatt |
| BPSV40-GAL4(del2-94)-EGFP_insert_R | aatctatctgtgacggcatctttattcacattatcttgtacaaataatccaacttttcttttctttt ttggagattc |
| Gal4(del615-767)-EGFP_insert | aatcgaatgctgagaataacgagaccgcacaattattacaacaaattaacgccaatt ttaatcaaagtgggaatattgctgatagctcattgtccttcac |
| Gal4(del768-881)-EGFP_insert | aacagctgcaatcattagtgccactgaccccgtctgctttgtttggtggcgccgctctag aactagtggatccgctgcaggaattcgatatcgtgtctaa |
| Gal4(del615-881)-EGFP_insert | aatcgaatgctgagaataacgagaccgcacaattattacaacaaattaacgccgctc tagaactagtggatccgctgcaggaattcgatatcgtgtctaa |
| GAL10_tet_NAT_F | ttttgttattgtacgtggggcagttgacgtcttatcatatgtcaaagtcatttgcgaagtctg cattgacaagttgctacacg |
| GAL10_tet_NAT_R | aaaagtttcaagacggcaatctctttttactgcatctcgtcagttggcaacttgccaaga ctgagtgagtacagtacgtgacg |
| tetTargetL_pTL541_F | ctgcattgacaagttgctacacg |

| | |
|---|---|
| tetTargetR_pTL541_R | ctgagtgagtacagtacgtgacg |
| 3'RNR2_tet_NAT_F | gagtatctctctatatatttctttttacgcagtctcttcaatctctttatctgcattgacaagttg ctacacg |
| 3'RNR2_tet_NAT_R | tatgattcagtgtatatatatatatataaataagaggtgcgaaagcccacctgagtgag tacagtacgtgacg |
| 5'RNR2_tet_NAT_F | gcataggaagccgaagtcgaacaagaagcaggcaaagtttagagcactgcctgc attgacaagttgctacacg |
| 5'RNR2_tet_NAT_R | gttgagaattttattttcctcctagtttttcttttttgagtgcggagggctgagtgagtacagt acgtgacg |
| insert_GAL4(del768-881) | aacagctgcaatcattagtgccactgaccccgtctgctttgtttggtggctaaaatgaat cgtagatactgaaaaaccccgcaagttcacttcaactgtg |
| GAL80::KANMX-F | tccttgccgaccagcgtatacaatctcgatagttggtttcccgttctttccactcccgtctat cgataccgtcgacctcg |
| GAL80::KANMX-R | ctcagtattcgtttttataacgttcgctgcactgggggccaagcacagggcaagatgctt ctcgactagtggatctgata |
| med15del_insert | tgccgtactcaaagatcaaggattaaaacgctatttcttttaaatctgctacatttgaagtt tccatacttttgatacttttgaagttacttcgtttggt |
| Gal4-GFP-BPSV40_insert_F | gtcttgttagaatttgttactgctgctggtattacccatggtatggatgaattgtacaaaaa aaggacagcagatggttc |
| Gal4-GFP-BPSV40_insert_R | atgtggggggagggcgtgaatgtaagcgtgacataactaattacatgactcgaccag ttaaacttttcttttctttttggagattca |
| Gal4-GFP-3V5_insert_F | gatggtccagtcttgttacc |
| Gal4-GFP-3V5_insert_R | gggagggcgtgaatgtaagcgtgacataactaattacatgactcgaccagttatggat ctgtactatccagtcc |

REFERENCES


1. Erdős, G., Pajkos, M., and Dosztányi, Z. (2021). IUPred3: prediction of protein disorder enhanced with unambiguous experimental annotation and visualization of evolutionary conservation. Nucleic Acids Res. *49*, W297–W303. 10.1093/nar/gkab408.

2. Erdős, G., and Dosztányi, Z. (2020). Analyzing Protein Disorder with IUPred2A. Curr. Protoc. Bioinforma. *70*, e99. 10.1002/cpbi.99.

3. Laughery, M.F., Hunter, T., Brown, A., Hoopes, J., Ostbye, T., Shumaker, T., and Wyrick, J.J. (2015). New vectors for simple and streamlined CRISPR-Cas9 genome editing in Saccharomyces cerevisiae. Yeast Chichester Engl. *32*, 711–720. 10.1002/yea.3098.

4. Brouwer, I., Patel, H.P., Meeussen, J.V.W., Pomp, W., and Lenstra, T.L. (2020). Single-Molecule Fluorescence Imaging in Living Saccharomyces cerevisiae Cells. STAR Protoc. *1*, 100142. 10.1016/j.xpro.2020.100142.

5. Donovan, B.T., Huynh, A., Ball, D.A., Patel, H.P., Poirier, M.G., Larson, D.R., Ferguson, M.L., and Lenstra, T.L. (2019). Live-cell imaging reveals the interplay between transcription factors, nucleosomes, and bursting. EMBO J. *38*. 10.15252/embj.2018100809.

6. Brouwer, I., Kerklingh, E., Leeuwen, F. van, and Lenstra, T.L. (2021). Dynamic epistasis analysis reveals how chromatin remodeling regulates transcriptional bursting. 2021.12.15.472793. 10.1101/2021.12.15.472793.

7. Edelstein, A.D., Tsuchida, M.A., Amodaj, N., Pinkard, H., Vale, R.D., and Stuurman, N. (2014). Advanced methods of microscope control using μManager software. J. Biol. Methods *1*, e10. 10.14440/jbm.2014.36.

8. Coulon, A., Ferguson, M.L., de Turris, V., Palangat, M., Chow, C.C., and Larson, D.R. (2014). Kinetic competition during the transcription cycle results in stochastic RNA processing. eLife *3*, e03939. 10.7554/eLife.03939.

9. MacKinnon, J.G. (2007). Bootstrap Hypothesis Testing. Work. Pap.

10. Schneider, C.A., Rasband, W.S., and Eliceiri, K.W. (2012). NIH Image to ImageJ: 25 years of image analysis. Nat. Methods *9*, 671–675. 10.1038/nmeth.2089.

11. Stael, S., Miller, L.P., Fernández-Fernández, Á.D., and Van Breusegem, F. (2022). Detection of Damage-Activated Metacaspase Activity by Western Blot in Plants. Methods Mol. Biol. Clifton NJ *2447*, 127–137. 10.1007/978-1-0716-2079-3_11.

12. Masuda, C.A., Xavier, M.A., Mattos, K.A., Galina, A., and Montero-Lomeli, M. (2001). Phosphoglucomutase is an in vivo lithium target in yeast. J. Biol. Chem. *276*, 37794–37801. 10.1074/jbc.M101451200.

13. Lenstra, T.L., Coulon, A., Chow, C.C., and Larson, D.R. (2015). Single-Molecule Imaging Reveals a Switch between Spurious and Functional ncRNA Transcription. Mol. Cell *60*, 597–610. 10.1016/j.molcel.2015.09.028.

14. Dovrat, D., Dahan, D., Sherman, S., Tsirkas, I., Elia, N., and Aharoni, A. (2018). A Live-Cell Imaging Approach for Measuring DNA Replication Rates. Cell Rep. *24*, 252–258. 10.1016/j.celrep.2018.06.018.
